# Intranasally Administered EVs from hiPSC-derived NSCs Alter the Transcriptomic Profile of Activated Microglia and Conserve Brain Function in an Alzheimer’s Model

**DOI:** 10.1101/2024.01.18.576313

**Authors:** Leelavathi N Madhu, Maheedhar Kodali, Raghavendra Upadhya, Shama Rao, Bing Shuai, Yogish Somayaji, Sahithi Attaluri, Maha Kirmani, Shreyan Gupta, Nathaniel Maness, Xiaolan Rao, James Cai, Ashok K. Shetty

## Abstract

Antiinflammatory extracellular vesicles (EVs) derived from human induced pluripotent stem cell (hiPSC)-derived neural stem cells (NSCs) hold promise as a disease-modifying biologic for Alzheimer’s disease (AD). This study directly addressed this issue by examining the effects of intranasal administrations of hiPSC-NSC-EVs to 3-month-old 5xFAD mice. The EVs were internalized by all microglia, which led to reduced expression of multiple genes associated with disease-associated microglia, inflammasome, and interferon-1 signaling. Furthermore, the effects of hiPSC-NSC-EVs persisted for two months post-treatment in the hippocampus, evident from reduced microglial clusters, inflammasome complexes, and expression of proteins and/or genes linked to the activation of inflammasomes, p38/mitogen-activated protein kinase, and interferon-1 signaling. The amyloid-beta (Aβ) plaques, Aβ-42, and phosphorylated-tau concentrations were also diminished, leading to better cognitive and mood function in 5xFAD mice. Thus, early intervention with hiPSC-NSC-EVs in AD may help maintain better brain function by restraining the progression of adverse neuroinflammatory signaling cascades.

## Introduction

The pathological changes associated with cognitive and mood impairments in Alzheimer’s disease (AD) include lasting neuroinflammation, extracellular deposition of amyloid-beta 42 (Aβ42) plaques, intraneuronal neurofibrillary tangles, synapse loss, hyperphosphorylated tau (p-tau), and neurodegeneration^1–4^. Deposition of Aβ plaques likely contributes to the progression of neuroinflammation, the loss of synapses, p-tau accumulation, and neurodegeneration^3^. Current therapeutic strategies for AD are not efficacious in slowing its progression^5^. Therefore, novel therapies proficient in restraining the progression of multiple pathological changes, including neuroinflammatory signaling cascades, and maintaining better cognitive and mood function for extended periods after the initial AD diagnosis have great significance^3^.

Transplantation of neural stem/progenitor cells (NSCs), derived from various sources, including the human induced pluripotent stem cells (hiPSCs) has improved function in several brain disease models^6–10^. However, multiple safety issues have hampered the clinical translation of hiPSC-derived NSC grafts^11–12^. Moreover, unlike other brain diseases, NSC grafting for AD is challenging as it involves pathological alterations in multiple brain regions^13–14,10^. Besides, the functional recovery after NSC grafting has been mainly attributed to the secretome of the NSC graft-derived cells^8,15^. Also, the vital component of the NSC secretome has been identified as extracellular vesicles (EVs), as they facilitate the transfer of genetic information and proteins from NSCs into host cells^15^. EVs, carrying a cargo of miRNAs and proteins from parental cells, can modify the function of recipient cells by transferring components^16–18^. Therefore, NSC-derived EVs, likely retaining most of the therapeutic effects of NSCs, can promote a cell-free therapy for AD^19–21^. Indeed, EVs derived from hiPSC-NSCs (hiPSC-NSC-EVs) have robust therapeutic properties because of antiinflammatory miRNAs and proteins carried by them^19,22^. Additionally, hiPSC-NSC-EVs are a much safer alternative to hiPSC-NSCs, as they do not replicate, and can readily cross the blood-brain barrier. EVs are also amenable for repeated, non-invasive intranasal (IN) dispensation as an allogeneic off-the-shelf product for treating neurodegenerative diseases because EVs can be stored frozen and used immediately after thawing without losing their biological activity. Furthermore, EVs quickly permeate the entire brain after an IN administration, including in an AD model^23–26^.

Therefore, using 3-month-old 5x familial AD (5xFAD) mice, we investigated whether IN administrations of hiPSC-NSC-EVs in the early stage of AD would moderate the activation of microglia, the associated neuroinflammatory signaling cascades, Aβ plaques, and p-tau, leading to better cognitive and mood function. These EVs, purified from hiPSC-NSC culture media using anion-exchange and size-exclusion chromatographic methods, naturally carry a cargo of antiinflammatory miRNAs and proteins^19,22^. Single cell-RNA-sequencing (scRNA-seq) revealed the proficiency of hiPSC-NSC-EVs to modulate multiple genes linked to activation of disease-associated microglia (DAM), NOD-, LRR- and pyrin domain-containing protein 3 (NLRP3)-inflammasomes, and interferon-1 (IFN-1) signaling in microglia. Such modulatory effects persisted even two months post-treatment in the hippocampus, perceptible from reduced concentrations of proteins and/or genes causing the activation of NLRP3 inflammasomes and p38/mitogen-activated protein kinase (p38/MAPK) signaling. Additionally, proteins involved in the cyclic GMP-AMP synthase and stimulator of interferon genes (cGAS-STING) pathway that activate interferon-1 signaling genes were also reduced in the hippocampus. These changes led to reduced Aβ, p-tau, and better cognitive and mood function.

## Results

### hiPSC-NSC-EVs displayed EV-specific markers and ultrastructure and were internalized by microglia in 5xFAD mouse brain following IN administration

The hiPSC-NSC-EVs were small EVs (30-250 nm in diameter, average size = 145 nm; Suppl Fig. 2 [A]). They expressed multiple EV-specific markers, including CD63, CD81, and Alix, and lacked cytoplasmic markers expressed in parental NSCs such as calnexin and cytochrome C (Suppl Fig. 1 [B]). Transmission electron microscopy revealed vesicles displaying double membrane (Suppl Fig. 1 [C]). The miRNA and protein composition of these hiPSC-NSC-EVs have been described in our previous study^19^. The ability of IN-administered EVs to target neurons and glia in all brain regions of the 5xFAD mice has been presented in our recent study^26^. Microglia in virtually all forebrain regions of 5xFAD mice, including plaque-associated microglia (PAM), internalized IN administered PKH-26+ EVs when examined 45 minutes post-administration (Suppl Fig. 2 [A-H]).

### hiPSC-NSC-EV incorporation triggered transcriptomic changes within microglia of 5xFAD mice

Supplementary Figure 3 outlines the experimental design employed in the study. First, we investigated transcriptomic changes in microglia induced by hiPSC-NSC-EVs at 72 hours post-administration using the scRNA seq. The pattern of t-SNE plot of microglia from the AD-Veh group differed from corresponding plots from naïve and AD-EVs groups (Fig. 1 [A]). The AD-Veh group showed upregulation of 8,735 genes and downregulation of 1,300 genes vis-à-vis the naïve group. In contrast, the AD-EVs group demonstrated upregulation of 4,050 genes and downregulation of 1,402 genes compared to the naïve group. Compared to the AD-Veh group, the AD-EVs group displayed upregulation of 1,506 genes and downregulation of 8,280 genes (Fig. 1 [B], Suppl Fig 4). Microglia with a high DAM gene signature were apparent in the AD-Veh group compared to the naïve group (Fig. 1 [C-D]). The upregulated genes comprised *ctsd, ctsb, ctsl, ctsz, axl, gpnmb, spp1, timp2, c3, igf1, lyz2, cybb, apoe, lpl, fth1, cst7, trem2, tyrobp, lilrb4a, itgax, cd9, cd63, cd74, csf1, ccl6*. Notably, most of these genes, except *gpnmb, c3, trem2, cd9* were downregulated in the AD-EVs group (Fig. 1 [C-D], Suppl Fig. 5 [A]). Furthermore, multiple genes linked to NLRP3 inflammasomes *(nfkb1, rela, nlrp3, pycard, casp1, il1b, il18)* were upregulated in the AD-Veh group. The expression of many of these genes *(rela, nlrp3, pycard, casp1, il1b)* was reduced in the AD-EVs group (Fig. 1 [E-F], Suppl Fig. 5 [B]). Moreover, the expression of microglial homeostatic genes *(p2ry12, p2ry13, cx3cr1, and cd33)* was downregulated in the AD-Veh group but normalized in the AD-EVs group (Fig. 1 [G-H]). Furthermore, the AD-Veh group displayed upregulation of multiple genes linked to the signaling of IFN-1 (18 genes), IFN-γ (27 genes), and IL-6 (23 genes). The AD-EVs group displayed normalized or reduced expression of many upregulated genes related to these signaling pathways. These include genes *b2m, bst2, gbp3, gbp7, h2-k1, ifitm3, lgals3bp, psmb8, psmb10, rnf2, stat1, stat2, tap2 and tapbp* (IFN-1 signaling genes), *cd86, cd274, fcgr1, hfe, jak2, psmb9, stat1* (IFN-γ signaling genes), and *cd14, crlf2, csf1, csf2rb, csf3r, ifnar1, ifngr2, il1b, il10rb, il13Ra1, il15ra, IL17ra, ltbr, myd88, ptpn2, socs3, stat1, stat2, stat3, tlr2, and tnf* (IL-6 signaling genes) (Extended Fig. 1 and Suppl Figs 6-8). Thus, IN-administration of hiPSC-NSC-EVs in 5xFAD mice resulted in significant transcriptomic changes implying suppression of NLRP, IFN-1, IFN-γ, and IL-6 signaling.

**Figure 1:**
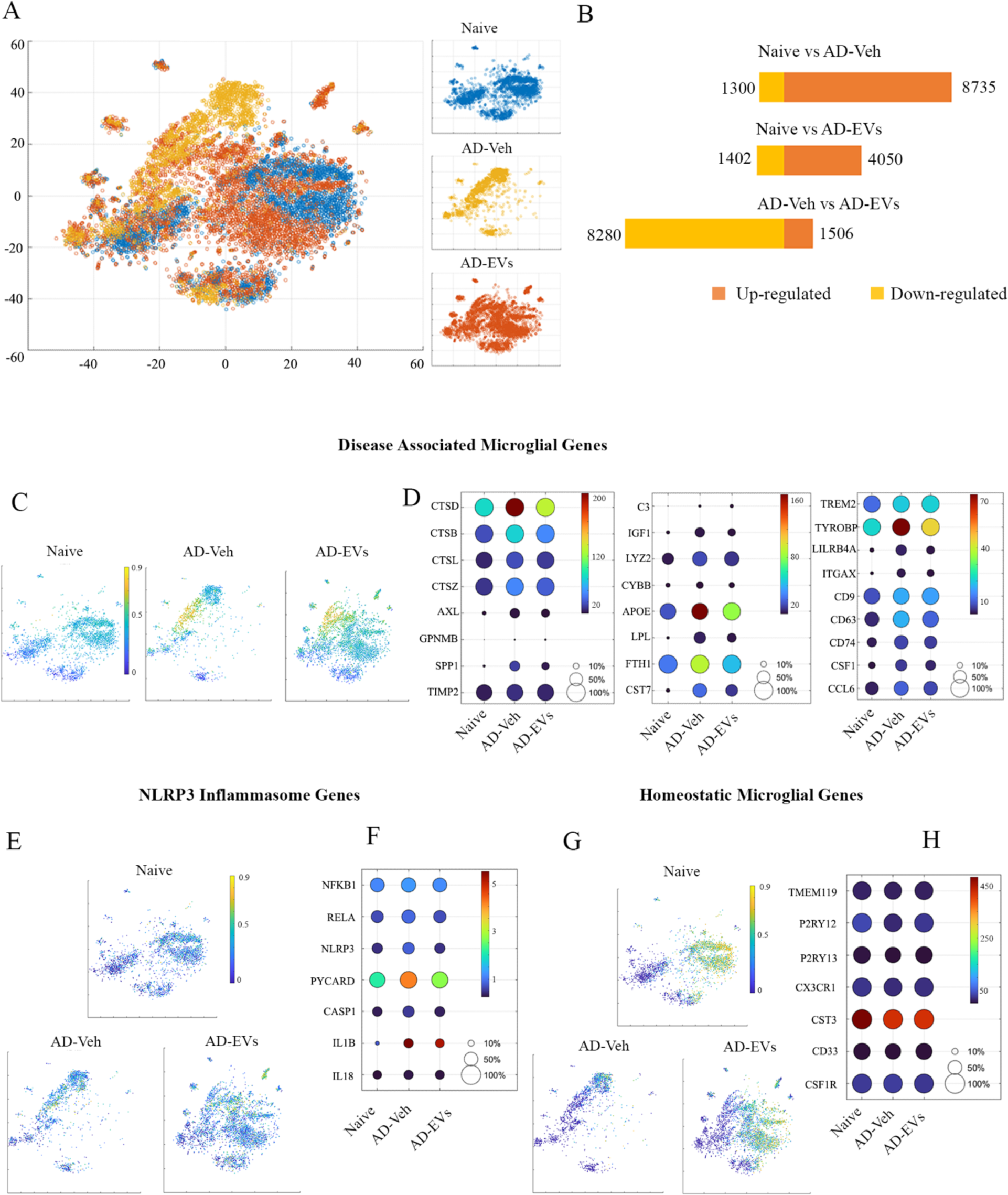
Intranasal administration of extracellular vesicles from human induced pluripotent stem cell-derived neural stem cells (hiPSC-NSC-EVs) altered the expression of genes linked to disease-associated microglia (DAM) and NOD-, LRR- and pyrin domain-containing protein 3 (NLRP3) inflammasome activation in 5xFAD mice microglia when observed 72 hours post-EVs administration. Figure A illustrates the distinct t-SNE plots of microglia in naïve, AD-Veh, and AD-EVs groups. B represents the total number of microglial genes upregulated and downregulated in the AD-Veh and AD-EVs groups compared to the naïve group. C and D represent the Ucell score scatter plot and dot plots of different DAM genes in microglial cells of naïve, AD-Veh, and AD-EVs groups. E and F represent the Ucell score scatter plot and dot plots of different NLRP3 inflammasome genes in microglial cells of different groups. G and H represent the Ucell score scatter plot and dot plots of different homeostatic microglial genes in microglial cells of different groups.

### hiPSC-NSC-EVs treatment reduced Aβ42-induced activation of cultured human iMicroglia

We next examined whether hiPSC-NSC-EVs are also proficient in moderating the expression of genes in cultured human iMicroglia challenged with Aβ42 oligomers. iMicroglia, generated from hiPSCs and expressing the microglia-specific transmembrane protein 119 (TMEM119) (Fig. 2 [A-E]), were exposed to Aβ42 (1 µM) for 24 hours. One set of cultures received 6 x 10^9^ hiPSC-NSC-EVs at 4 hours after Aβ42 exposure (Fig. 2 [F]). The expression of homeostatic microglia genes *(tmem119, p2ry12)* did not differ between naïve, Aβ42-exposed, and Aβ42-exposed and hiPSC-NSC-EV treated iMicroglia (p>0.05; Fig. 2 [G-H]). However, compared to the naïve iMicroglia, the expression of multiple genes linked to microglia activation *(cd68, cx3cr1, c1qa)*, DAM *(ctsd, apoe, lpl, fth1)* and inflammation *(il1b, tnfa)* were upregulated in the Aβ42-exposed iMicroglia (p<0.05-0.01, Fig. 2 [I-R]) but not in the Aβ42-exposed iMicroglia treated with hiPSC-NSC-EVs (p>0.05, Fig. 2 [I-R]). However, investigation of phagocytosis function revealed that hiPSC-NSC-EVs mediated reduced activation of iMicroglia did not interfere with their phagocytosis function (Suppl Fig. 9). Thus, hiPSC-NSC-EVs can also moderate human microglial activation without compromising their phagocytosis function.

**Figure 2:**
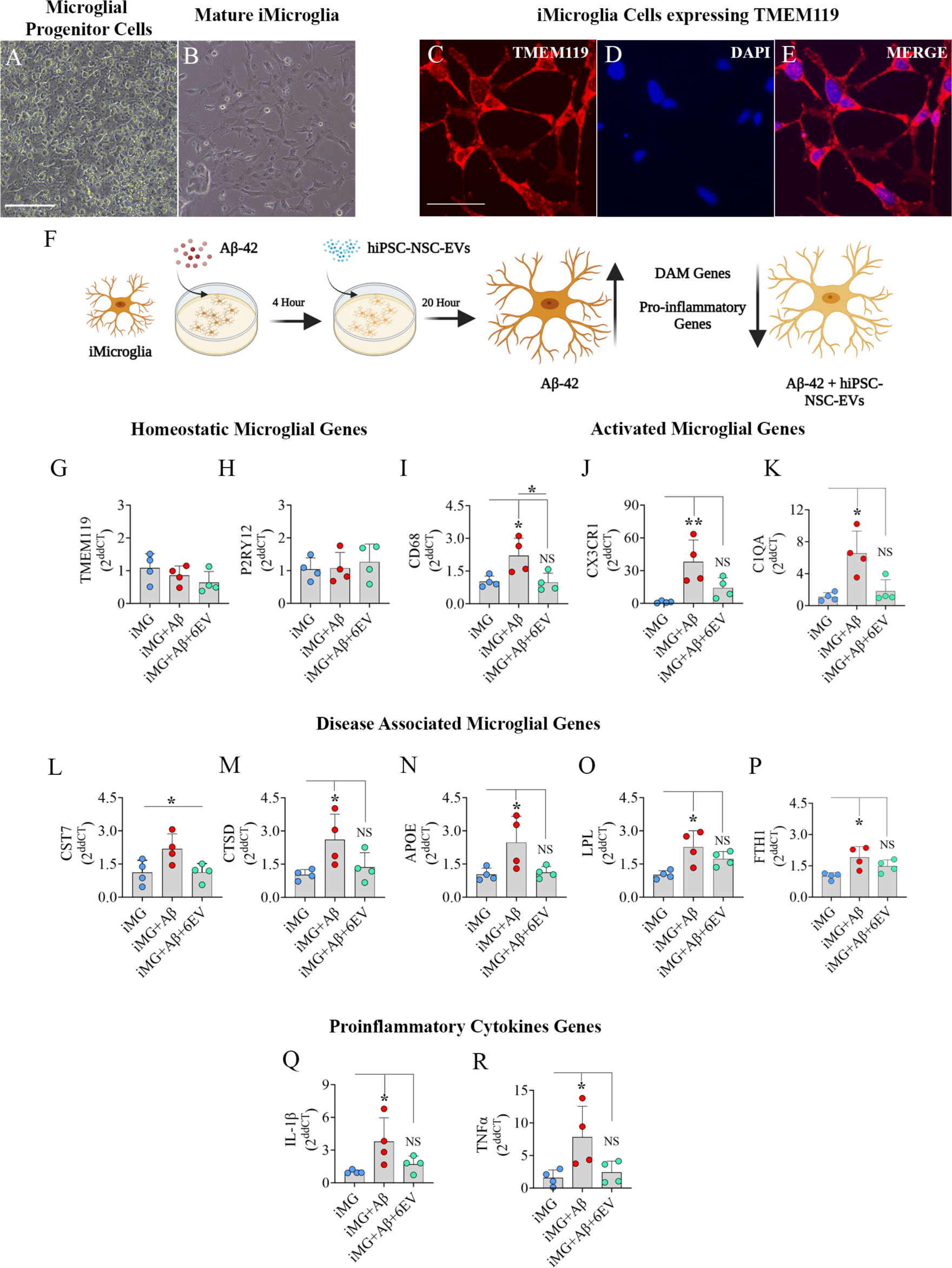
Intranasal administration of extracellular vesicles from human induced pluripotent stem cell-derived neural stem cells (hiPSC-NSC-EVs) suppressed Aβ42-induced activation of iMicroglia derived from hiPSCs. Images A and B show the progenitor and mature microglia from hiPSCs. C-D images confirm TMEM119 expression in mature microglia. F is a diagrammatic representation of the experiment involving Aβ42 exposure to iMicroglia followed by hiPSC-NSC-EV treatment. Bar charts G-H, I-K, L-P, and Q-R respectively compare the expression of homeostatic genes (G-H), activated microglia genes (I-K), disease-associated microglia (DAM) genes (L-P), and proinflammatory cytokine genes (Q-R) in iMicroglia between control, Aβ42-exposed, and Aβ42-exposed and hiPSC-NSC-EVs treated cultures. Scale bar, A-E= 100 µm; *, p<0.05; **, p<0.01; NS, not significant.

### hiPSC-NSC-EV administration preserved cognitive, memory and mood function at 5 months of age

We first measured the competence of animals to recognize minor changes in the immediate environment, a hippocampus-dependent cognitive function, using an object location test (OLT; Fig. 3 [A]). The majority of the animals (10-13 males and 6-13 females) met the criteria employed for this task (i.e., exploration of objects ≥20 seconds in trial-2 [T2]). Proficiency for object location memory was intact in both males and females in the age-matched naïve control group when analyzed separately or together (p<0.01-0.0001; Fig. 3 [B, F, J]), but was impaired in both males and females in the AD-Veh group (p>0.05, Fig. 3 [C, G, K]). Contrastingly, both males and females in the AD-EVs group displayed no cognitive impairment (p<0.05-0.001, Fig. 3 [D, H, L]). Intact cognitive function in naive and AD-EV groups was evident from the propensity of animals to explore the object in novel place (OINP) for longer durations than the object in familiar place (OIFP). Analysis of the OINP-DI values using a one-sample t-test revealed proficiency for the cognitive task in the naïve group (males and females, alone or together, p<0.05-0.01) and the AD-EVs group (males, p<0.05, both sexes, p<0.01) but not in the AD-Veh group (Fig. 3 [E, I, M]). The results were not influenced by variable object exploratory behavior between groups, as the total object exploration times (TOETs) in T2 did not differ between groups (p>0.05, Suppl Fig. 10) [A-B]).

**Figure 3:**
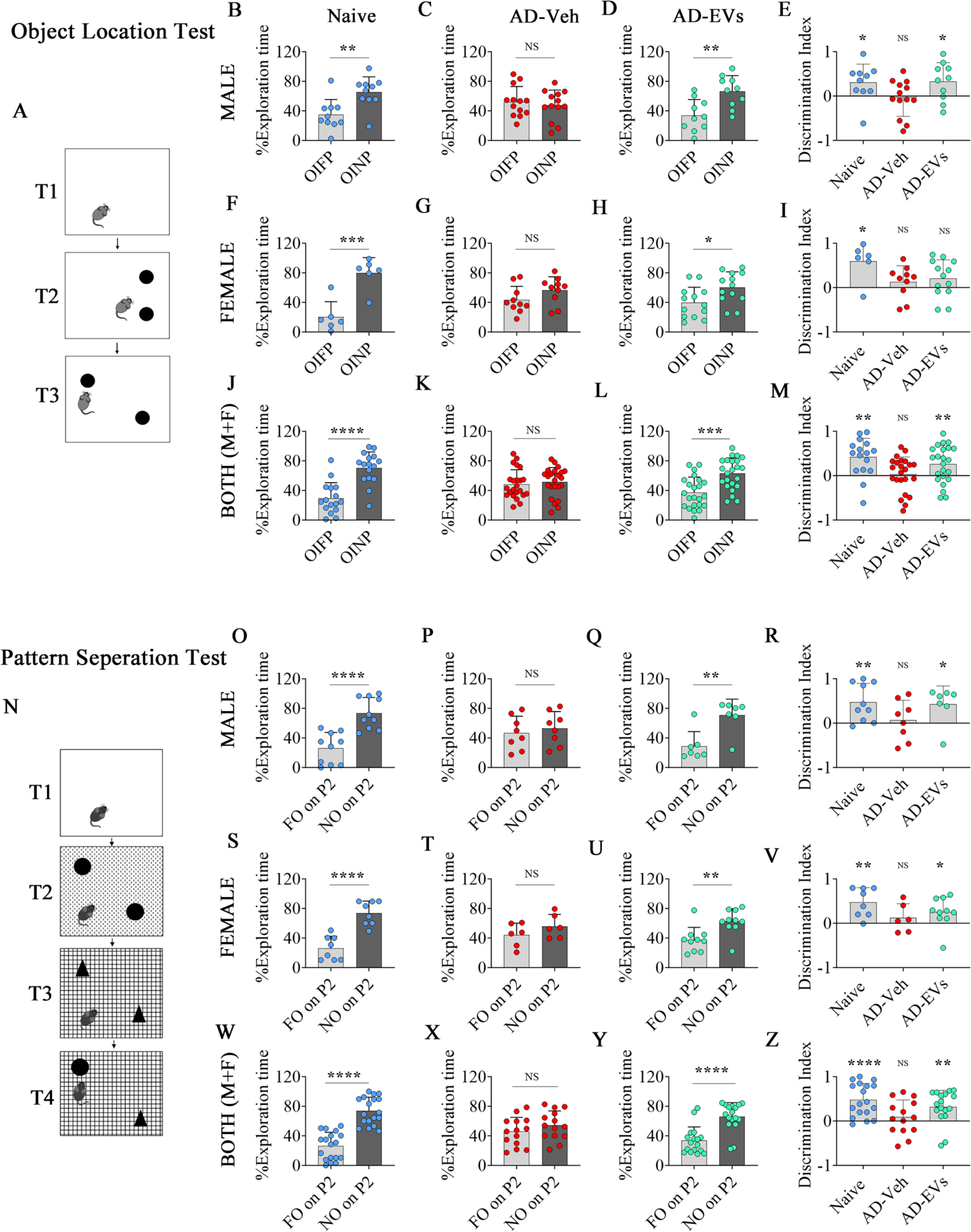
Intranasal administration of extracellular vesicles from human induced pluripotent stem cell-derived neural stem cells (hiPSC-NSC-EVs) to 5xFAD mice preserved cognitive function. Cartoon A depicts different trials (T1-T3) in an object location test (OLT). The bar charts in B-D, F-H, J-L compare percentages of object exploration times spent with the object in the familiar place (OIFP) vis-à-vis the object in the novel place (OINP) in naïve (B, F, J), AD-Veh (C, G, K) and AD-EVs (D, H, L) in males (B-D), females (F-H) and when both males and females were considered together (J-L). The bar charts E, I, M compare the OINP discrimination index (DI) for males (E), females (I), and both sexes (M) using a one-sample t-test. Cartoon N depicts different trials (T1-T4) in a pattern separation test (PST). The bar charts in O-Q, S-U, W-Y compare percentages of object exploration times spent with the familiar object on pattern 2 (FO on P2) vis-à-vis the novel object on P2 (NO on P2) in naïve (O, S, W), AD-Veh (P, T, X) and AD-EVs (Q, U, Y) in males (O-Q), females (S-U) when both males and females were considered together (W-Y) together. The bar charts R, V, Z compare the DI for the NO on P2 for males (R), females (V), and both sexes (Z) using a one-sample t-test. *, p<0.05; **, p<0.01; ***, p<0.001; and ****, p<0.0001; NS, not significant.

Next, we determined the pattern separation ability of animals using a pattern separation test (PST; Fig. 3 [N]), which examined the proficiency for discriminating the novel object (NO) from the familiar object (FO) kept on a second-floor pattern (P2) in T4 of PST. Most animals (9-11 males and 9-12 females) in every group met the criteria (i.e., exploration of objects ≥20 seconds in T2 and T3). Adeptness for recognizing the NO on P2 in T4 was intact in both males and females in the naïve (p<0.0001, Fig. 3 [O, S, W]) and AD-EVs (p<0.01-0.0001, Fig. 3 [Q, U, Y]) groups but impaired in the AD-Veh group (p>0.05, Fig. 3 [P, T, X]). Proficiency for pattern separation in naive and AD-EV groups was evident from the propensity of animals to explore the NO on P2 for longer durations than the FO on P2. Analysis of the NO on P2-DI values using a one-sample t-test confirmed competence for pattern separation in males and females of the naïve group (p<0.01-0.0001) and the AD-EVs group (p<0.05-0.01) but not in the AD-Veh group (Fig. 3 [R, V, Z]). As observed in OLT, the findings were not prompted by varying object exploratory behavior between groups, as the TOETs in T2 and T3 did not differ between groups (p>0.05, Suppl Fig. 10 [C-F].

Furthermore, an investigation of mood function using a sucrose preference test (SPT) (n=10-11 males and 9-10 females per group, Fig. 4 [A]) revealed anhedonia in the AD-Veh group compared to the naïve group but not in the AD-EVs group (Suppl Fig. 10 [G-L]). Absence of anhedonia in both sexes in naïve and AD-EV groups was evident from the preference of animals to drink higher amounts of sucrose-containing water than standard water (p<0.0001, Suppl Fig. 10 [G, I, J, L). In contrast, the presence of anhedonia in both sexes in the AD-Veh group was apparent from the lack of preference of animals to drink sweet water (p>0.05, Suppl Fig. 10 [H, K]). Comparison of the sucrose preference rate (SPR) across groups confirmed differences between naive and AD-Veh groups, and between AD-Veh and AD-EVs groups, for both males and females or when both sexes were considered together (p<0.01-0.0001, Fig 4 [B, C, D]). Thus, hiPSC-NSC-EVs intervention at three months of age in male and female 5xFAD mice prevented the occurrence of anhedonia at five months of age.

**Figure 4:**
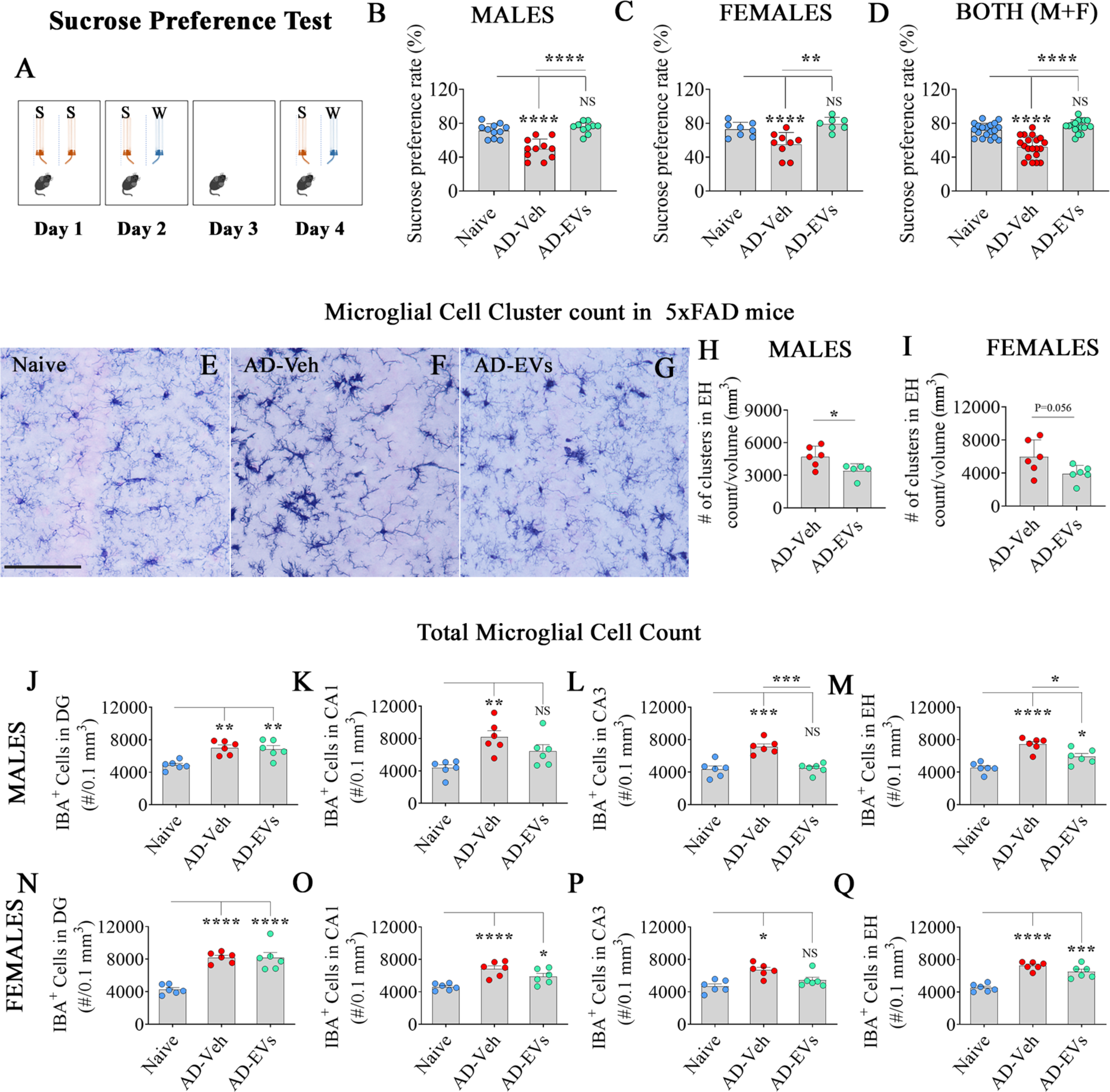
Intranasal administration of extracellular vesicles from human induced pluripotent stem cell-derived neural stem cells (hiPSC-NSC-EVs) to 5xFAD mice prevented anhedonia and reduced microglial clusters and numbers. Cartoon A depicts the experimental design employed for the sucrose preference test. The bar charts B-D compare the sucrose preference rate (SPR) in males (B), females (C) and males + females (D) across naïve, AD-Veh and AD-EVs groups. **, p<0.01; ****, p<0.0001; NS, not significant. Figure E-G illustrates the representative images of microglial clusters in naïve (E), AD-Veh (F) and AD-EVs (G) groups. Bar charts H-I compare the number of microglial clusters per mm^3^ unit area of the hippocampus in males (H) and females (I) between naive, AD-Veh, and AD-EV groups. Bar charts J-Q compare numbers of microglia in males (J-M) and females (N-Q) per 0.1 mm^3^ area of the dentate gyrus (J, N), CA1 subfield (K, O), CA3 subfield (L, P), and the entire hippocampus (M, Q) between naive, AD-Veh, and AD-EVs groups. Scale bar, E-G = 100 μm *, p<0.05; **, p<0.01; ***, p<0.001; ****, p<0.0001; NS, not significant.

### hiPSC-NSC-EVs administration reduced microglial clusters and numbers in the hippocampus

We quantified the number of microglial clusters in different hippocampal subfields (Fig. 4 [E-I]). IBA-1+ microglial clusters, were conspicuous in all hippocampal subfields in AD-Veh and AD-EVs groups. However, males in the AD-EVs group displayed a reduced number of microglial clusters per cubic millimeter (mm^3^) compared to the AD-Veh group (p<0.05, Fig.4 [H]). The females in the AD-EVs group also showed a similar trend, but the difference was not significant (p=0.056, Fig. 4 [I]). Furthermore, in both sexes, compared to the naive group, the AD-Veh group displayed an increased number of microglia in all subfields and the entire hippocampus (p<0.05-0.0001, Fig. 4 [J-Q]). Males in the AD-EVs group exhibited reduced numbers of microglia in the CA3 subfield and the entire hippocampus compared to the AD-Veh group (p<0.05-0.001; Fig. 4 [L, M]). However, females in the AD-EVs group displayed similar numbers of microglia as the AD-Veh group in all hippocampal subfields (Fig 4 [N-Q]). Thus, hiPSC-NSC-EV treatment reduced microglial clusters and numbers in male 5xFAD mice. The females showed a similar trend, but the reductions were not statistically significant.

### hiPSC-NSC-EVs treatment maintained lower expression of DAM genes for extended periods

We quantified and compared the expression of multiple DAM genes in the hippocampus at 5 months of age (i.e., ∼2 months post-EV treatment) across groups. Compared to the naïve control group, the AD-Veh group displayed increased expression of *cst7, spp1, lpl, apoe, fth1, and tyrobp* in males (p<0.05-0.0001, Fig. 5 [A-F]) and *cst7, lpl, fth1, tyrobp, and ctsd* in females (p<0.05-0.001, Fig. 5 [I, K, M-N, P]). However, the expression of most of these genes in both males and females in the AD-EVs group was normalized to levels in the naïve group (p>0.05, Fig. 5 [B-F, I, K, M, N, P]). The expression of some genes was also reduced in the AD-EVs group compared to the AD-Veh group (p<0.05, Fig. 5 [A, C, D, I]). Thus, the modulatory effects of hiPSC-NSC-EVs treatment on DAM genes observed at 72 hours post-EV administration persisted at ∼2 months post-EV treatment.

**Figure 5:**
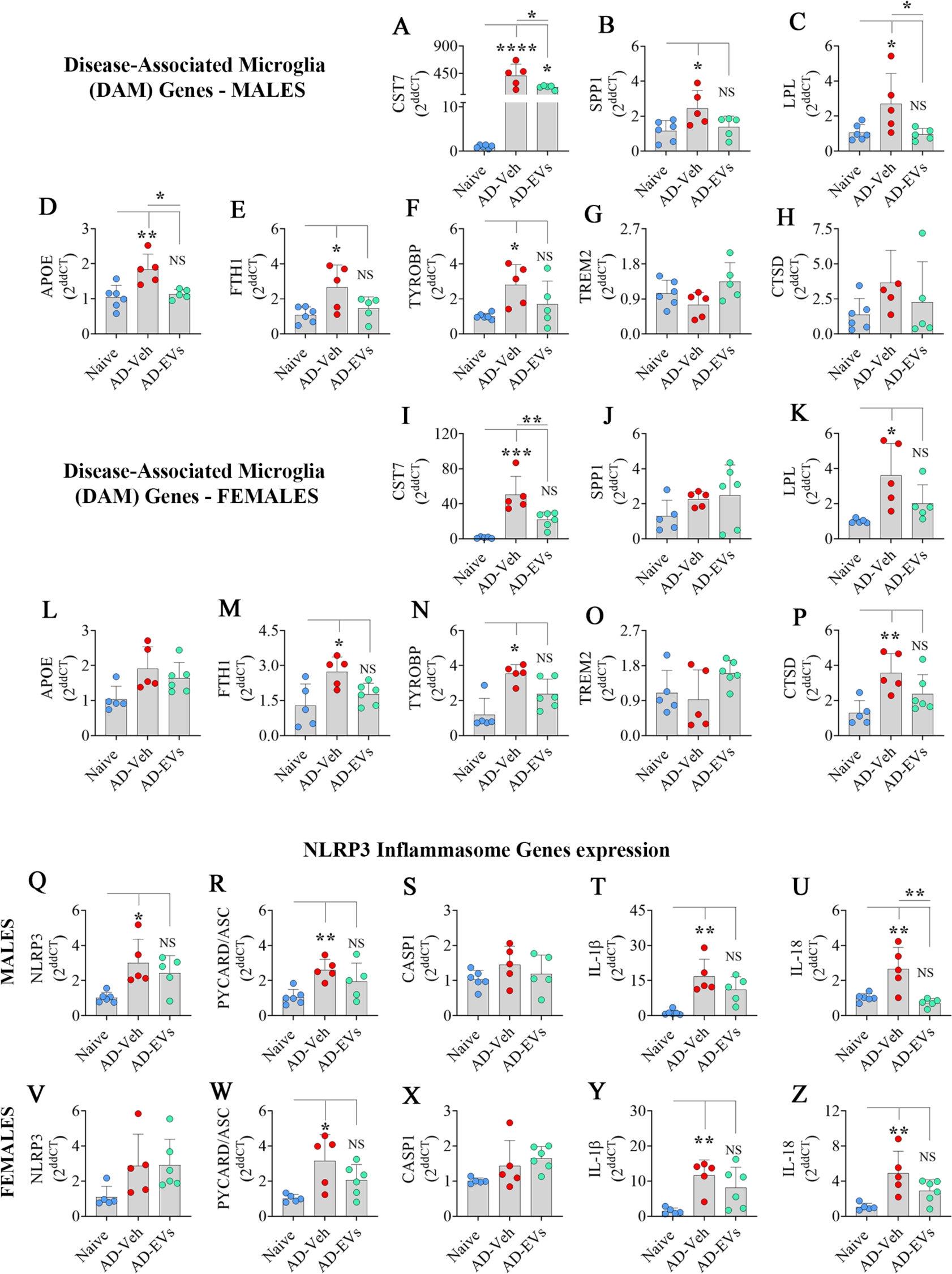
Intranasal administration of extracellular vesicles from human induced pluripotent stem cell-derived neural stem cells (hiPSC-NSC-EVs) to 5xFAD mice normalized the expression of genes linked to disease associated microglia (DAM) and inflammasome activation. The bar charts A-P compare the expression of DAM genes (cst7, spp1, lpl, apoe, fth1, tyrobp, trem2, ctsd) in the hippocampus of males (A-H) and females (I-P) between naïve, AD-Veh and AD-EVs groups. The bar charts Q-Z compare the expression of genes linked to NOD-, LRR- and pyrin domain-containing protein 3 (NLRP3) inflammasome activation (nlrp3, pycard, casp1, il-1β, il-18) in hippocampus of males (Q-U) and females (V-Z) between naïve, AD-Veh and AD-EVs groups. *, p<0.05; **, p<0.01; ***, p<0.001; ****, p<0.0001; NS, not significant.

### hiPSC-NSC-EVs administration induced enduring inhibition of NLRP3 inflammasome activation

We first measured and compared the expression of genes linked to NLRP3 inflammasome activation across groups. The AD-Veh group displayed increased expression of *nlrp3, pycard/asc, il1b and il18* genes in males (p<0.05-0.01; Fig. 5 [Q-R, T-U]) and *pycard/asc, il1b,* and *il18* genes in females (p<0.05-0.01; Fig 5 [W, Y, Z]). However, in the AD-EVs group, the expressions of these genes were normalized to levels in the naïve group (p>0.05, Fig. 5 [Q-R, T-U, W, Y, Z]. Next, we quantified the extent of NLRP3 inflammasome complexes within microglia in the hippocampus through triple immunofluorescence for IBA-1, NLRP3, and ASC. Examples of microglia displaying NLRP3 inflammasome complexes (i.e., structures co-expressing NLRP3 and apoptosis-associated speck-like protein containing a caspase recruitment domain (CARD); ASC) are illustrated (Fig. 6 [A-I]). Compared to the naïve group, both males and females in the AD-Veh group displayed increased percentages of microglia presenting inflammasome complexes (p<0.01, Fig. 6 [J, K]). On the other hand, percentages of microglia containing inflammasome complexes in both sexes of the AD-EVs group matched percentages in the naïve group (p>0.05, Fig. 6 [J, K]) and were less than their counterparts in the AD-EVs group (p<0.05, Fig. 6 [J, K]). To confirm these further, we quantified the concentrations of NLRP3 inflammasome activation mediators (NFkB, NLRP3, ASC, and cleaved caspase-1) and end products (IL-1β and IL-18) in the hippocampus. Compared to the naïve group, both sexes in the AD-Veh group displayed increased concentrations of all these proteins (p<0.05-0.0001, Fig. 6 [L-Q, R-W]). Notably, the concentrations of all these proteins were either normalized to levels in the naïve group (p>0.05, Fig. 6 [M-O, Q, R-W]) or went below the level of the naïve group (Fig. 6 [L, P]) in males and females in the AD-EVs group. Thus, the repressing effects of hiPSC-NSC-EVs on NLRP3 inflammasome genes observed at 72 hours post-EV administration persisted at ∼2 months post-EV treatment.

**Figure 6:**
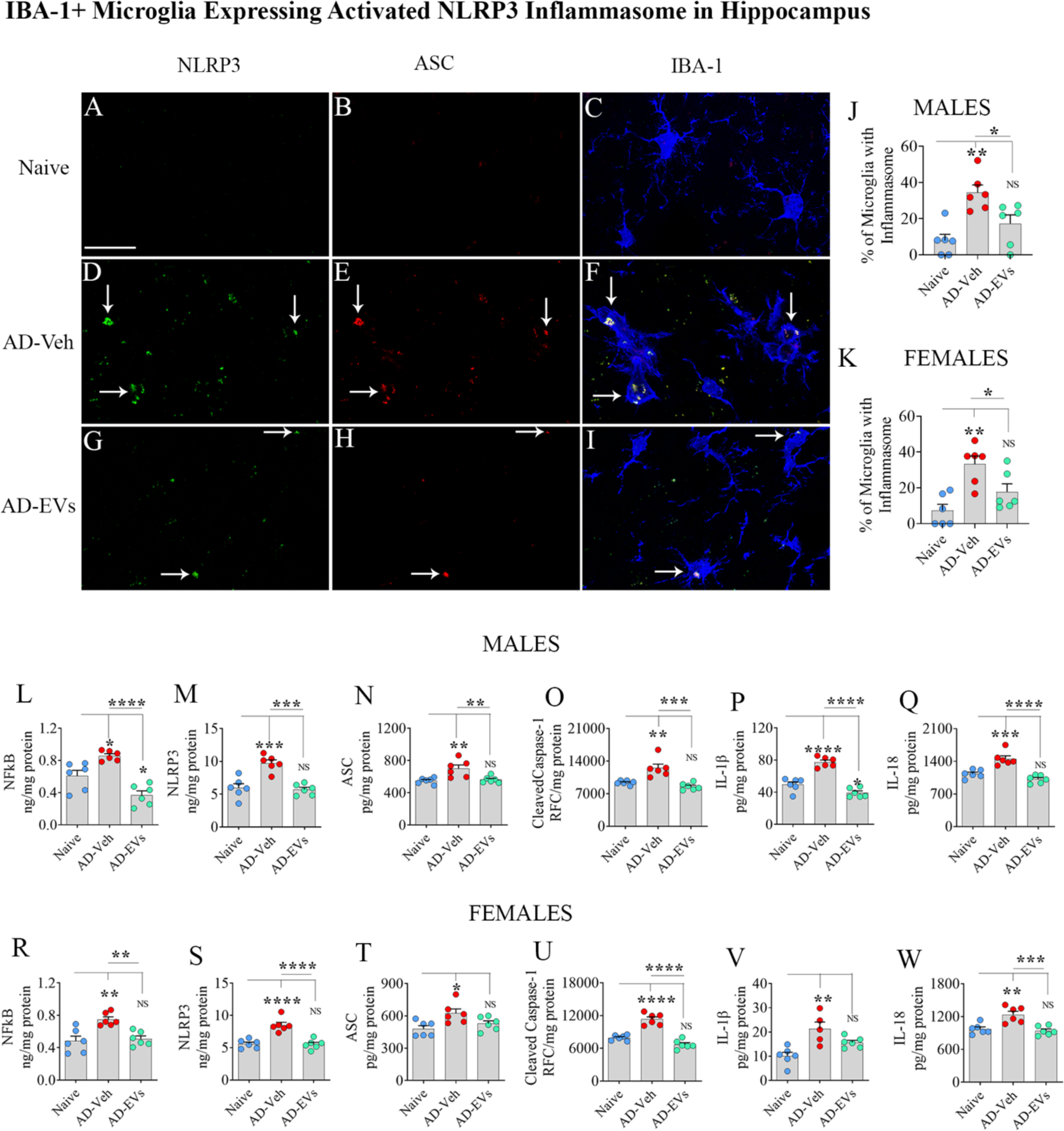
Intranasal administration of extracellular vesicles from human induced pluripotent stem cell-derived neural stem cells (hiPSC-NSC-EVs) to 5xFAD mice inhibited NOD-, LRR- and pyrin domain-containing protein 3 (NLRP3) inflammasome complex formation and activation. Figures A-I illustrate examples of NLRP3 inflammasome complexes co-expressing NLRP3 (green) and apoptosis-associated speck-like protein containing a CARD (ASC, red) in IBA-1+ microglia (blue) from the CA3 subfield of the hippocampus in male mice from naïve, AD-Veh and AD-EVs groups. The bar charts J-K compare the percentages of microglia with inflammasome complexes in males (J) and females (K). The bar charts L-W compare the concentrations of mediators of NLRP3 inflammasome activation (NF-kB, NLRP3, ASC, and cleaved caspase-1; L-O, male and R-U, female) and end products (IL-1β, IL-18; P-Q, and V-W) in males (L-Q) and females (R-W) between naive, AD-Veh, and AD-EVs groups. Scale bar, A-I = 25 μm; *, p<0.05; **, p<0.01; ***, p<0.001; ****, p<0.0001; NS, not significant.

### hiPSC-NSC-EVs treatment prevented hyperactivation of p38/MAPK signaling

Increased release of proinflammatory cytokines IL-18 and IL-1β following NLRP3 inflammasome activation leads to downstream p38/MAPK hyperactivation in acute and chronic inflammatory conditions^24^. Therefore, we first compared the concentrations of different p38/MAPK signaling components, including myeloid differentiation primary response 88 (MyD88), a small GTPase rat sarcoma virus (Ras), phospho-p38 MAPK, and activator protein-1 (AP-1) in naïve, AD-Veh, and AD-EVs groups. Next, we compared the concentrations of some known end products of p38/MAPK hyperactivation, including IL-6, tumor necrosis factor -alpha (TNF-α), and IL-8 and macrophage inflammatory protein-1 alpha (Mip-1α). Compared to the naïve group, males in the AD-Veh group displayed elevated levels of MyD88, Ras, p38/MAPK, and AP-1 levels (p<0.05-0.01; Fig. 7 [A-D]). Females in the AD-Veh group also showed a similar trend, but only Ras and p38/MAPK increases were statistically significant (p<0.05-0.01; Fig. 7 [J, K]). Notably, in the AD-EVs group, the concentrations of these proteins were normalized to levels in the naive control group (p>0.05, Fig. 7 [A-D, J, K]). Furthermore, in both males and females, the concentrations of IL-8, TNF-α, and Mip-1α were elevated in the AD-Veh group but not in the AD-EVs group (p<0.05-0.001; Fig. 7 [F-H, N-P]). Females in the AD-Veh group, in addition, showed upregulation of IL-6 compared to the naive group (p<0.01; Fig. 7 [M]), which was normalized in the AD-EVs group (p>0.05, Fig. 7 [M]). Thus, inhibition of NLRP3 inflammasome activation mediated by hNSC-EVs prevented the downstream hyperactivation of p38/MAPK signaling.

**Figure 7:**
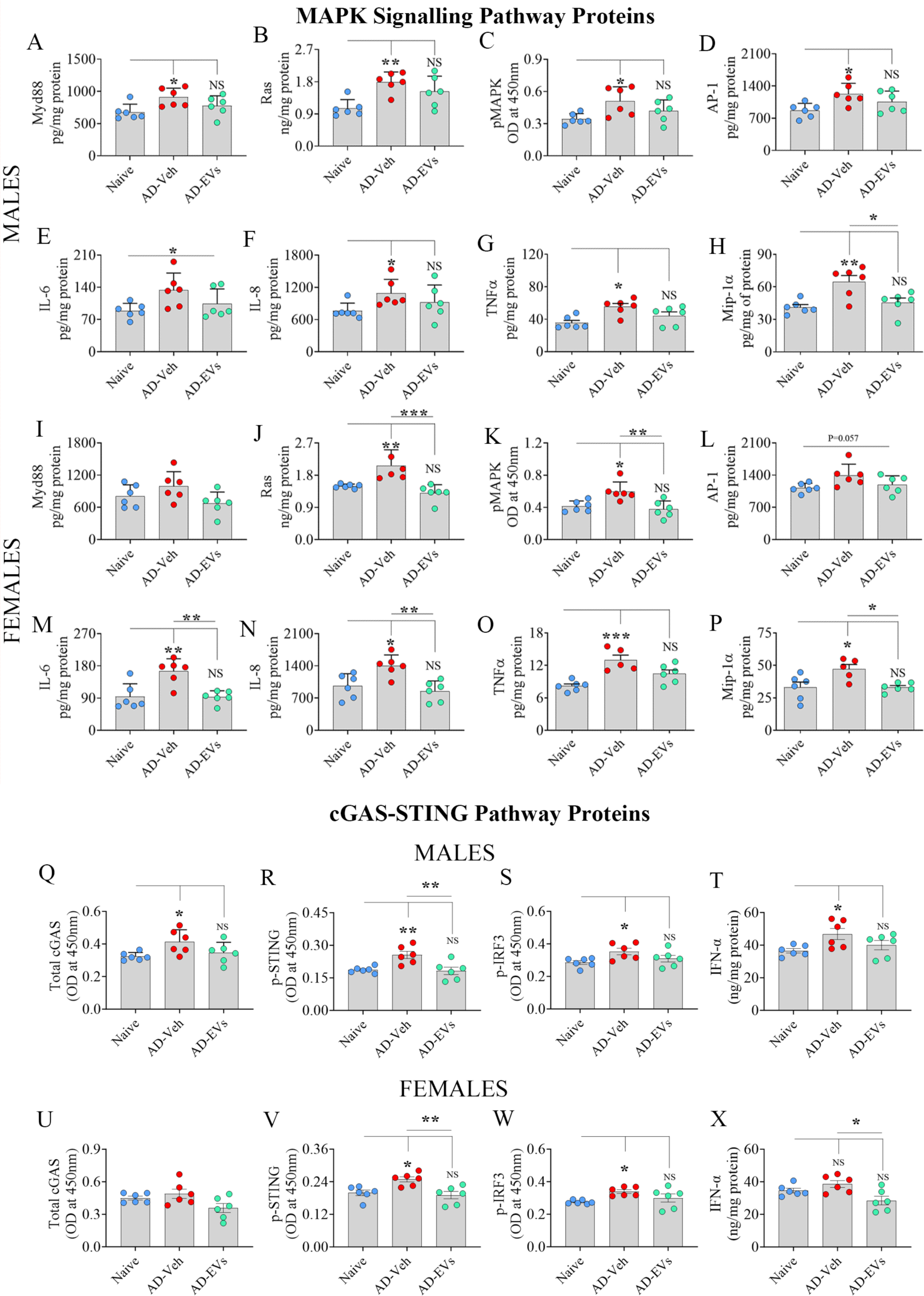
Intranasal administration of extracellular vesicles from human induced pluripotent stem cell-derived neural stem cells (hiPSC-NSC-EVs) to 5xFAD mice thwarted the activation of p38 mitogen-activated protein kinase and cyclic GMP-AMP synthase (cGAS), and phosphorylated stimulator of interferon genes (p-STING) signaling. The bar charts A-P compare the concentrations of various components of p38/MAPK activation (MyD88, Ras, pMAPK, AP-1; A-D and I-L) and end products (IL-6, IL-8, TNFα, Mip-1α; E-H and M-P) in males (A-H) and females (I-P) between naive, AD-Veh, and AD-EVs groups. *, p<0.05; **, p<0.01; ***, p<0.001; NS, not significant. The bar charts Q-X compare total cGAS (Q, U), p-STING (R, V), p-IRF3 (S, W), IFN-α (T, X) across groups in males (Q-T) and females (U-X). *, p<0.05; **, p<0.01; ***, p<0.001; NS, not significant.

### hiPSC-NSC-EVs administration induced enduring inhibition of cGAS-STING signaling

We quantified the concentrations of various proteins causing the activation of the cGAS-STING pathway, an upstream event activating IFN-1 signaling in AD and 5xFAD mice^27^. Compared to the naive group, the hippocampus from the AD-EVs group displayed increased concentrations of total cGAS, p-STING, phosphorylated interferon regulatory factor 3 (p-IRF3), and IFN-α in males (p<0.05-0.01, Fig. 7 [Q-T]), and p-STING and p-IRF3 in females (p<0.05, Fig. 7 [V-W]). In contrast, in the AD-EVs group, the concentrations of these proteins in both sexes did not differ from the naive group (p>0.05, Fig. 7 [Q-X]). Thus, the modulatory effects of hiPSC-NSC-EVs on IFN-1 signaling observed at 72 hours post-EV administration persisted at ∼2 months post-EV treatment.

### hiPSC-NSC-EVs administration reduced amyloid plaque load and p-tau

Comparing area fractions of brain tissue covered by amyloid plaques in the hippocampus revealed that males and females in the AD-EVs group displayed significantly lower plaque load than the AD-Veh group. (p<0.05, Extended Fig. 2 [A-D]). Also, the hippocampal concentrations of soluble Aβ42 were reduced in males and females of the AD-EVs group compared to the AD-Veh group (p<0.05-0.01; Extended Fig. 2 [E-F]). Furthermore, the hippocampal concentrations of p-tau were reduced in males and females of the AD-EVs group compared to the AD-Veh group (p<0.05-0.01; Extended Fig. 2 [G-H]). Thus, hiPSC-NSC-EV-mediated reductions in neuroinflammatory cascades led to reduced amyloid plaques, Aβ42 production, and p-tau in the 5xFAD mouse brain.

## Discussion

The results of this study, in male and female 5xFAD mouse models, provide novel evidence that IN administrations of hiPSC-NSC-EVs in the early stage of AD could slow down the progression of neuroinflammation, and the extent of Aβ plaques and p-tau in the hippocampus and thereby maintain better cognitive and mood function at an advanced stage of the disease. 5xFAD mice received hiPSC-NSC-EVs at three months of age, an age at which these mice start to display Aβ plaques, microgliosis, and a proinflammatory milieu in the brain^28–29^, and interrogated with behavioral tests at ∼4.5 months of age, a stage where untreated 5xFAD mice display cognitive and mood impairments^30–31^. The neuropathology was assessed at five months of age, a timepoint in which 5xFAD mice exhibited increased microglial activation with upregulation of multiple DAM genes, and activation of NLRP3 inflammasomes, and the p38/MAPK and cGAS-STING-IFN-1 signaling cascades. Remarkably, male and female 5xFAD mice receiving hiPSC-NSC-EVs displayed better cognitive function and no anhedonia at 4.5 months of age compared to their counterparts receiving the vehicle. Moreover, hiPSC-NSC-EVs treatment led to reductions in the expression of genes and/or proteins linked to DAM, NLRP3, p38/MAPK and cGAS-STING-IFN-1 hyperactivation, Aβ plaques, Aβ42 and p-tau concentrations at 5 months of age.

The concept of using hiPSC-NSC-EVs to restrain the neuroinflammatory cascades involved in AD pathogenesis stemmed from our earlier studies demonstrating the proficiency of these EVs to mediate robust antiinflammatory properties. For example, hiPSC-NSC-EVs have shown proficiency to suppress IL-6 release from lipopolysaccharide (LPS)-stimulated mouse macrophages, and IL-1β and TNF-α release from LPS-stimulated human iMicroglia^19,22^. They also thwarted neuroinflammation after a brain insult such as status epilepticus^19^ or LPS-induced peripheral inflammation^21^. Furthermore, these EVs are inherently enriched with antiinflammatory miRNAs and proteins, evidenced by small RNA sequencing and proteomic studies^19^. Our previous protein knockdown and miRNA inhibition studies have identified miR-21-5p and pentraxin-3 (PTX3) as the key antiinflammatory molecules in these EVs^22^. Such findings are consistent with the ability of miR-21-5p to regulate NF-kB, enhance IL-10, and inhibit TNFα release^32–34^ and the proficiency of PTX3 to enhance the neuroprotective type 2 astrocytes, and regulate the entry of peripheral immune cells into the brain^35–36^. hiPSC-NSC-EVs also carry miR-103a, proficient in reducing neuroinflammation via prostaglandin-endoperoxide synthase-2 inhibition^37^, hemopexin capable of transforming proinflammatory microglia into antiinflammatory phenotypes^38^, and galectin-3 binding protein (Gal-3BP) competent in diminishing the NF-kB signaling^39^.

The entry of hiPSC-NSC-EVs into microglia in all brain regions led to reduced expression of multiple genes linked to DAM, NLRP3 inflammasome activation, and IFN-1 signaling. Such changes implying the modulation of activated microglia have implications because microglia, though recognize and remove a significant amount of Aβ in the early stage of AD, undergo hyperactivation, perpetuate chronic neuroinflammation and contribute to disease progression and dementia in the advanced stage^40–42^. Such transformed microglia lose homeostatic molecules and functions^43^ and acquire DAM or “neurodegenerative” phenotype^44^. DAM is typified by downregulation of ‘‘homeostatic’’ microglial genes *p2ry12, p2ry13, cx3cr1, cd33, and tmem119*^45^ and upregulation of genes involved in lysosomal, phagocytic, and lipid metabolism pathways such as *apoe, ctsd, lpl, tyrobp, and trem2*^46–47^. Notably, hiPSC-NSC-EVs-mediated transformation of microglia involved increased expression of homeostatic genes and reduced expression of DAM genes, implying the proficiency of hiPSC-NSC-EVs to transform DAM into a less inflammatory microglia. Testing the effects of hiPSC-NSC-EVs on human iMicroglia exposed to Aβ oligomers also confirmed the competence of hiPSC-NSC-EVs to downregulate DAM gene expression in activated human iMicroglia.

hiPSC-NSC-EVs treatment normalized/reduced the expression of many genes linked to the NLRP3 inflammasome activation. Similar effects were also seen in human iMicroglia exposed to Aβ oligomers. Activation of NLRP3 inflammasomes within microglia in AD occurs as a response to Aβ, as Aβ fibrils can induce IL-1β release from glia through NLRP3 inflammasome activation, and Aβ oligomers and fibrils can directly interact with NLRP3 inflammasome components and induce its activation^48–50^. Additionally, p-tau-derived paired helical filament-6 (PHF6) peptides can induce NLRP3 inflammasome activation^51^. Inhibition of NLRP3 inflammasome activation is beneficial because of its contribution to AD pathogenesis, particularly its ability to perpetuate chronic neuroinflammation via downstream hyperactivation of p38/MAPK signaling, increase tau phosphorylation, and induce cognitive dysfunction. Such concept is supported by findings of increased NLRP3 inflammasome end products (IL-1β and IL-18) in AD patients^52–53^, and better cognitive function, reduced chronic neuroinflammation and Aβ42 in AD models with the inhibition of NLRP3 inflammasome activation^54–56^. Aβ induced tau pathology is also linked to NLRP3 inflammasome activation, as NLRP3 knockout in tauopathy mice reduced tau phosphorylation by regulating tau kinases and phosphatases^57^.

IN administration of hiPSC-NSC-EVs also normalized/reduced the expression of many genes linked to IFN-1, IFN-γ, and IL-6 signaling within microglia in 5xFAD mice. Type-I IFNs, including IFN-α and IFN-β, bind to the IFN-1 receptor complex and activate signaling via kinases JAK1 and TYK2, leading to STAT1 and STAT2 phosphorylation, which results in the upregulation of thousands of interferon-stimulated genes (ISGs)^58^. Type-I IFNs are chronically activated within dysfunctional microglia in AD^59^, and IFNα/β transcripts and ISGs are upregulated in the brains of AD patients^60–61^. Also, the expression of IRF7, a transcription factor controlling IFN-1 signaling correlates with AD progression^60–61^. Furthermore, both amyloid and tau models of AD display an increased population of IFN-responsive microglia overexpressing ISGs^62–63^. While type-I IFNs protect against infection, their overproduction in neurodegenerative conditions can cause adverse effects, including increased synapse loss, microglial activation, and aggregation of Aβ42 and p-tau^64–65,61^. On the other hand, increased IFN-γ signaling in microglia can transform them into neurotoxic phenotypes capable of impairing neural network rhythms and cognitive functions, and causing neurodegeneration^66^. Additionally, increased IL-6 can contribute to memory impairment in AD^67^. Thus, the adeptness of hiPSC-NSC-EVs to reduce IFN-1, IFN-γ and IL-6 signaling within activated microglia in 5xFAD mice is advantageous for slowing AD progression.

Furthermore, transcriptomic changes induced by hiPSC-NSC-EVs on microglia in 5xFAD mice at 72 hours post-EV administration persisted at two months post-EV treatment in the hippocampus. This was evidenced by male and female 5xFAD mice receiving hiPSC-NSC-EVs displaying reductions in a) microglial clusters and numbers, b) the expression of multiple genes linked to DAM and NLRP3 inflammasome activation, c) percentages of microglia presenting NLRP3 inflammasomes, d) the concentrations of NLRP3 inflammasome activation mediators and end products, e) proteins involved in p38/MAPK hyperactivation, and f) multiple proinflammatory cytokines. Reduced clusters of microglia suggest reductions in PAM as such clusters are seen around Aβ plaques^68^, which is likely a consequence of diminished plaque density with hiPSC-NSC-EVs treatment. Reduced numbers of microglia imply reduced proliferation of microglia, whereas the reduced percentages of microglia displaying NLRP3 inflammasome complexes indicate reduced microglial activation^69^. Such changes in microglia were associated with reductions in the expression of genes linked to DAM and NLRP3 inflammasomes and the concentration of proteins linked to NLRP3 inflammasome activation (NF-kB, NLRP3, ASC, cleaved caspase-1 and IL-1β and IL-18). Dampened NLRP3 inflammasome activation led to reduced secretion of IL-1β and IL-18, which prevented the hyperactivation of p38/MAPK signaling, Such an effect was evidenced by reduced concentrations of MyD88, Ras, pMAPK, and AP-1, IL-6, IL-8, TNF-α, and Mip-1α. Preventing the hyperactivation of p38/MAPK signaling in AD is beneficial because p38/MAPK promotes NF-kB activation, tau phosphorylation, and glutamate excitotoxicity, impairs synaptic plasticity and autophagy and promotes neurodegeneration^70^. Moreover, knockdown of p38/MAPK has been shown to alleviate microglia activation and postpone cognitive decline in animal models of AD^71–72^. Additionally, at ∼2 months post-EV treatment, animals in the AD-EVs group displayed diminished cGAS-STING activation implicated in IFN-1 signaling^27^.

In both AD models and patients, Aβ plaques increase progressively and are surrounded by activated microglia with DAM genes signature and an impaired ability for phagocytosis and Aβ clearance^73–74^. Since hiPSC-NSC-EVs treatment transformed microglia into less inflammatory states, evident from the reduced expression of DAM and NLRP3 inflammasome genes and diminished clustering^46–47^, it is likely that the proficiency of microglia for Aβ phagocytosis and clearance improved. Such a conclusion is supported by diminished microglial clusters around smaller and fewer Aβ plaques in the AD-EVs group. Decreased formation of Aβ plaques following hiPSC-NSC-EVs treatment likely involved two mechanisms. First, since activated microglia spread Aβ seeds to form new plaques^75–76^, modulation of activated microglia by hiPSC-NSC-EVs likely diminished such seeding. Second, hiPSC-NSC-EVs are naturally enriched with Gal-3BP, which can suppress Aβ production by inhibiting β-secretase^39^. Such an effect is supported by the reduced concentration of Aβ42 in the hippocampus of the AD-EVs group. On the other hand, reduced p-tau in the AD-EVs group is a consequence of diminished NLRP3 inflammasome activation, as Aβ induced tau pathology is linked to the extent of NLRP3 inflammasome activation^57^.

In conclusion, IN administrations of hiPSC-NSC-EVs in 5xFAD mice result in their incorporation into microglia, which alters microglial transcriptomic signature and leads to sustained reductions in downstream neuroinflammatory signaling cascades and better cognitive and mood function. The results also imply that microglia-mediated chronic neuroinflammation in AD contributes profoundly to the progression of cognitive and mood function decline, and EV-based biologics capable of modulating their overactivation without affecting their homeostatic and Aβ clearing functions can considerably slow down the progression of AD pathogenesis.

## Methods

### Culturing of hiPSC-derived NSCs, purification, and characterization of hiPSC-NSC-EVs

Protocols for generating NSCs from hiPSCs (IMR90-4; Wisconsin International Stem Cell Bank, Madison, WI, United States), culturing of hiPSC-NSCs, isolating hiPSC-NSC-EVs using anion-exchange and size-exclusion chromatographic methods, characterizing EVs for various markers and ultrastructure are detailed in our previous studies^19,22^ and the supplementary file.

### Animals and study design

Both transgenic 5xFAD mice and their background (wild type) strain (B6SJLF1/J) were purchased from Jackson Labs (Cat No: 34840-JAX and 100012-JAX, Bar Harbor, Maine, USA) and maintained in our laboratory on B6/SJL genetic background by crossing 5xFAD transgenic male mice with B6SJLF1/J female mice. The animal care and use committee of Texas A&M University approved all studies conducted in this investigation. Two cohorts of 3-month-old 5xFAD mice were randomly assigned to vehicle (Veh) or hiPSC-NSC-EVs groups (AD-Veh group, n=26, 14 males and 12 females; AD-EVs group, n=27, 13 males and 14 females). A group of age-matched naïve control animals (n=24, 12 males and 12 females) served as the naïve group for neurobehavioral, immunohistochemical, and molecular studies. Furthermore, a separate set of animals (n=2/group, males) were employed for the initial scRNA-seq study. Additionally, four AD mice (males) were used to assess the incorporation of IN-administered PKH26-labeled hiPSC-NSC-EVs into microglia in different brain regions.

### Administration of hiPSC-NSC-EVs

Animals in AD-EVs and AD-Veh groups received IN administration of hiPSC-NSC-EVs or vehicle (sterile phosphate buffered saline, PBS). In each animal, following mild anesthesia, both nostrils were first treated with 10 µL of hyaluronidase (100 U; H3506; Sigma-Aldrich, St. Louis, MO, USA) in PBS to enhance the permeability of the nasal mucous membrane. Thirty minutes later, each mouse was gently held with the ventral side up, and the head was facing downward for the IN administration of EVs or PBS. EVs were suspended in PBS at a concentration of 30 x 10^9^/100μL and dispensed into both nostrils in 5-µL spurts separated by 5 minutes. The animals received two doses of 30 x 10^9^ EVs or Veh with a one-week interval between doses. An additional smaller cohort (n=4/group) of AD mice receiving 4 x 10^9^ EVs was euthanized 45 minutes after IN administration to confirm the distribution of EVs into microglia, including PAM in the hippocampus. The 5xFAD mice employed for the scRNA-seq study also received two doses of 30 x 10^9^ EVs or Veh (n=2/group), as described above.

### Isolation of live microglia from 5xFAD and naïve control mice and scRNA-seq studies

The 5xFAD mice employed for the scRNA-seq study were euthanized, and fresh brains were harvested 72 hours after the last dose of EV/Veh administration and processed immediately for the isolation of live microglia. The microglia were also isolated from the age-matched naïve control mice (n=2). The brain tissues from two animals were pooled in every group and processed for live single-cell suspension preparation using gentleMACS™ Tissue Dissociator (Miltenyi Biotec, Gaithersburg, MD, USA). The cell suspensions were incubated with CD11b microbeads (Miltenyi Biotec) and subjected to magnetic cell isolation through MACS® Separators to isolate microglia. Following live cell analysis and quality control measures, ∼5,000 live microglia were subjected to the scRNA-seq at the Texas A&M Institute for Genome Sciences and Society facility. Individually barcoded libraries were pooled and sequenced on a NextSeq mid-output paired-end sequencing run at 2×75 using NextSeq 500/550 Mid-Output v2.5 kit (Illumina, San Diego, CA, USA) according to the manufacturer’s instructions. The reads of scRNA libraries were aligned to the human GRCh38.p13 reference genome using Cell Ranger (version 7.0), and the resulting expression matrices were analyzed using scGEAToolbox^77^.

### Analysis of hiPSC-NSC-EVs effects on cultured hiPSC-derived microglia exposed to Aβ42 oligomers

A method described in a published protocol was employed to generate microglia from hiPSCs^78^. These microglia are referred to as iMicroglia from here onwards, and additional details on iMicroglia are available in the supplementary file. Three sets of mature iMicroglia were cultured on a Petri plate (n=8), of which two sets were exposed to Aβ42 oligomers at a concentration of 1 µM. Four hours after adding Aβ42 oligomers, one set of cultures received hiPSC-NSC-EVs (6 x 10^9^ EVs). Twenty hours later, iMicroglia from all culture sets were, dissociated, collected, washed, and processed for total RNA isolation and quantitative real-time PCR (qPCR) studies. We performed another set of experiments to understand potential changes in the phagocytic ability of iMicroglia after Aβ42 exposure with or without EV treatment. Twenty-three hours after Aβ42 exposure, fluorescent yellow-green latex beads (Sigma-Aldrich) were added to iMicroglia cultures treated with or without EVs. One hour later, images were captured, and the percentages of iMicroglia incorporating latex beads were quantified.

### Analyses of cognitive and mood function using neurobehavioral tests

Both male and female mice in naïve, AD-Veh, and AD-EVs groups were investigated with two neurobehavioral tests to measure cognitive function. The behavioral tests commenced three weeks after Veh/EVs administration and continued for a month (i.e., in the 5^th^ month of life). An OLT ascertained the proficiency of animals to discern minor changes in their immediate environment, a cognitive function entirely dependent on the hippocampus^79^. A PST ascertained the competence of animals to distinguish similar but not identical experiences in a nonoverlapping manner^80–81^, a measure of pattern separation function dependent on the dentate gyrus (DG) function. A SPT assessed mood function by examining anhedonia or depressive-like behavior. The detailed protocols employed for OLT, PST, and SPT are available in our previous reports^13,23–24,82^, and the supplementary file.

### Harvesting of brain tissues for immunohistochemical and molecular assays

The animals in all groups were euthanized following neurobehavioral tests when they were five months old. Fixed brain tissues were obtained for immunohistochemical studies, and fresh brain tissues were harvested for molecular assays. The details on the harvesting and processing of fixed and fresh tissues for immunohistochemical and molecular analyses are available in our published studies^21,82–85^ and the supplementary file.

### Quantification of numbers and clusters of microglia and Aβ42 plaques

The numbers of microglia were quantified via stereological counting of IBA1+ cells through the entire hippocampus in serial sections (every 20^th^ section) from naïve, AD-Veh, and AD-EVs groups (n=6/group), as described in our previous reports^86^. We also quantified the number of microglial clusters per cubic millimeter (mm^3^) volume from different hippocampus subfields. Image J was employed to quantify the extent of Aβ plaques in the hippocampus.

### Isolation of RNA from iMicroglia and mouse hippocampus for qPCR studies

Total RNA isolation kits were employed to isolate RNA from cultured iMicroglia (n=4/group; System Bioscience, Palo Alto, CA, USA) and the mouse hippocampal tissues (n=5-6/group; Qiagen, Germantown, MD, USA). The samples of the RNA (500ng/µL) were converted into cDNA using RT2 First Strand Kit (Qiagen). The qRT-PCR was performed using RT² SYBR Green qPCR Mastermix and Primer mix (GeneCopoeia, Rockville, Maryland, USA) for measuring human microglial gene expression in iMicroglia *(tmem119, p2ry12, cd68, cx3cr1, c1qa, cst7, ctsd, apoe, lpl, fth1, il1b, tnfa)* and mouse microglial gene expression in the hippocampus *(cst7, spp1, lpl, apoe, fth1, tyrobp, trem2, ctsd, nlrp3, pycard, casp1, il1b, il18)*.

### Measurement of NLRP3 inflammasome complexes in microglia

The brain tissue sections were processed for triple immunofluorescence staining and Z-section analysis using a Nikon confocal microscope or Leica THUNDER 3D Imager for measuring the percentages of microglia displaying NLRP3 inflammasome complexes (i.e., cells positive for IBA-1, NLRP3, and ASC) in the hippocampus^21^. All measurements were performed in the CA3 subfield of the hippocampus (3 sections/animal, n=6/group). The methods and antibodies employed for these studies are described in the supplementary file.

### Quantification of mediators and end products of NLRP3 inflammasome activation

The mediators and proinflammatory end products of NLRP3 inflammasome activation were quantified from hippocampal tissue lysates using ELISA (n=6/group), as detailed in our previous study^85^. Commercially available kits for measuring nuclear factor kappa B (NF-kB, Aviva Systems Biology, San Diego, CA, USA), NLRP3 (Abcam, Cambridge, MA, USA), ASC (MyBioSource, San Diego, CA, USA), cleaved caspase-1 (BioVision Inc, Milpitas, CA, USA), interleukin-18 (IL-18), IL-1β (R&D Systems, Minneapolis, MN, USA) were utilized.

### Measurement of proteins linked to hyperactivation of p38/MAPK and interferon signaling cascade

The hippocampal tissue lysates were utilized for quantifying the various markers involved in IL-18-mediated hyperactivation of the p38/MAPK signaling pathway and its proinflammatory end products (n=6/group). We followed the manufacturer’s protocol provided in individual ELISA kits for the MyD88 (Aviva Systems Biology), Ras (MyBioSource), p38 MAPK (Cell Signaling, Danvers, MA, USA), AP-1 (Novus Biologicals, Centennial, CO; USA), IL-6, TNF-α (R&D systems), IL-8 (Biomatik, Wilmington, DE, USA) and Mip-1α (Signosis, Santa Clara, CA, USA). The concentrations of these individual proteins were normalized to the 1 mg of total protein in hippocampal lysates, and values were compared across groups. Additionally, we also measured cGAS, p-STING, p-IRF3, and IFN-α concentrations in the hippocampus.

### Quantification of Aβ42 and p-tau concentrations

The five FAD mutations in 5xFAD mice lead to Aβ42 overproduction and increased Ser396 tau phosphorylation^87^. Therefore, we measured the concentrations of Aβ42 and p-tau in the hippocampal tissue lysates of naïve, AD-Veh, and AD-EVs groups. We followed the manufacturer’s protocol described in ELISA kits for measuring Aβ42 (Invitrogen, Waltham, MA, USA) and p-tau (Cell Signaling, Danvers, MA, USA). The concentrations of Aβ42 and p-tau were normalized to mg of protein in hippocampal lysates.

### Statistical analyses

Statistical analyses of data utilized Prism software 10.1. Within-group comparisons in neurobehavioral tests or two-group comparisons utilized a two-tailed, unpaired Student’s t-test or Mann-Whitney U-test for two datasets with significantly different standard deviations. All comparisons involving three or more datasets were analyzed using one-way ANOVA with Tukey’s multiple comparisons post-hoc tests. However, when individual groups did not pass the normality test Shapiro-Wilk test, Kruskal-Wallis test with Dunn’s post hoc tests were performed. The discrimination index (DI) values in neurobehavioral tests were compared to the hypothetical mean (zero) using a one-sample t-test. In all comparisons, p<0.05 was considered a statistically significant value.

## Data Availability

Source data are included with this original research article. Any additional data requests are available from the corresponding author upon request.

## Acknowledgments

The TEM images of hiPSC-NSC-EVs were taken at the Image Analysis Laboratory, Texas A&M Veterinary Medicine & Biomedical Sciences (RRIS: SCR_0222479).

## Authors’ contributions

Concept: AKS. Research design: AKS, LNM, MK, RU, SR, and JC. Data collection, analysis, and interpretation: LNM, MK, RU, SR, BS, YS, SA, MK, SG, NM, XR, JC, and AKS. Preparation of figure composites: LNM, MK, SR and AKS. Manuscript writing: LNM and AKS. All authors provided feedback, edits, and additions to the manuscript text and approved the final version of the manuscript.

## Ethical Approval

The animal care and experimental procedures were conducted per the animal protocol approved by the Animal Care and the Use Committee (IACUC) of Texas A&M University School of Medicine.

## Figure Legends

**Extended Figure 1:**
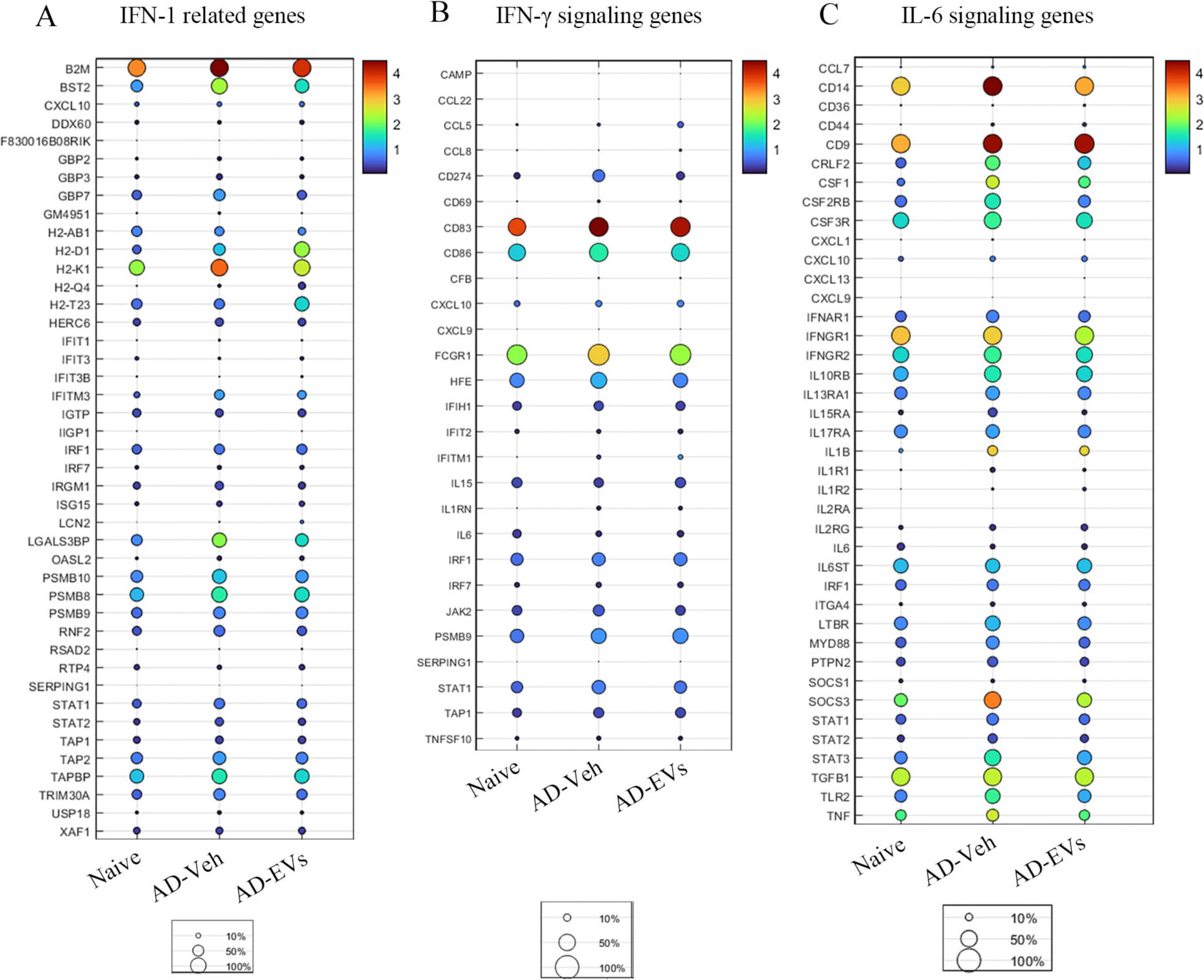
Intranasal administration of extracellular vesicles from human induced pluripotent stem cell-derived neural stem cells (hiPSC-NS-EVs) altered the expression of genes linked to interferon-1 (IFN-1), interferon-gamma (IFN-γ) interleukin-6 (IL-6) signaling in 5xFAD mice microglia when observed 72 hours post-EVs administration. Dot plots in A-C compare the expression of multiple genes linked to IFN-1, IFN-γ signaling, and IL-6 signaling pathways between naive, AD-Veh, and AD-EVs groups. The expression of most genes was upregulated in the AD-Veh group compared to the naive group but reduced in the AD-EVs treatment group compared to the AD-Veh group.

**Extended Figure 2:**
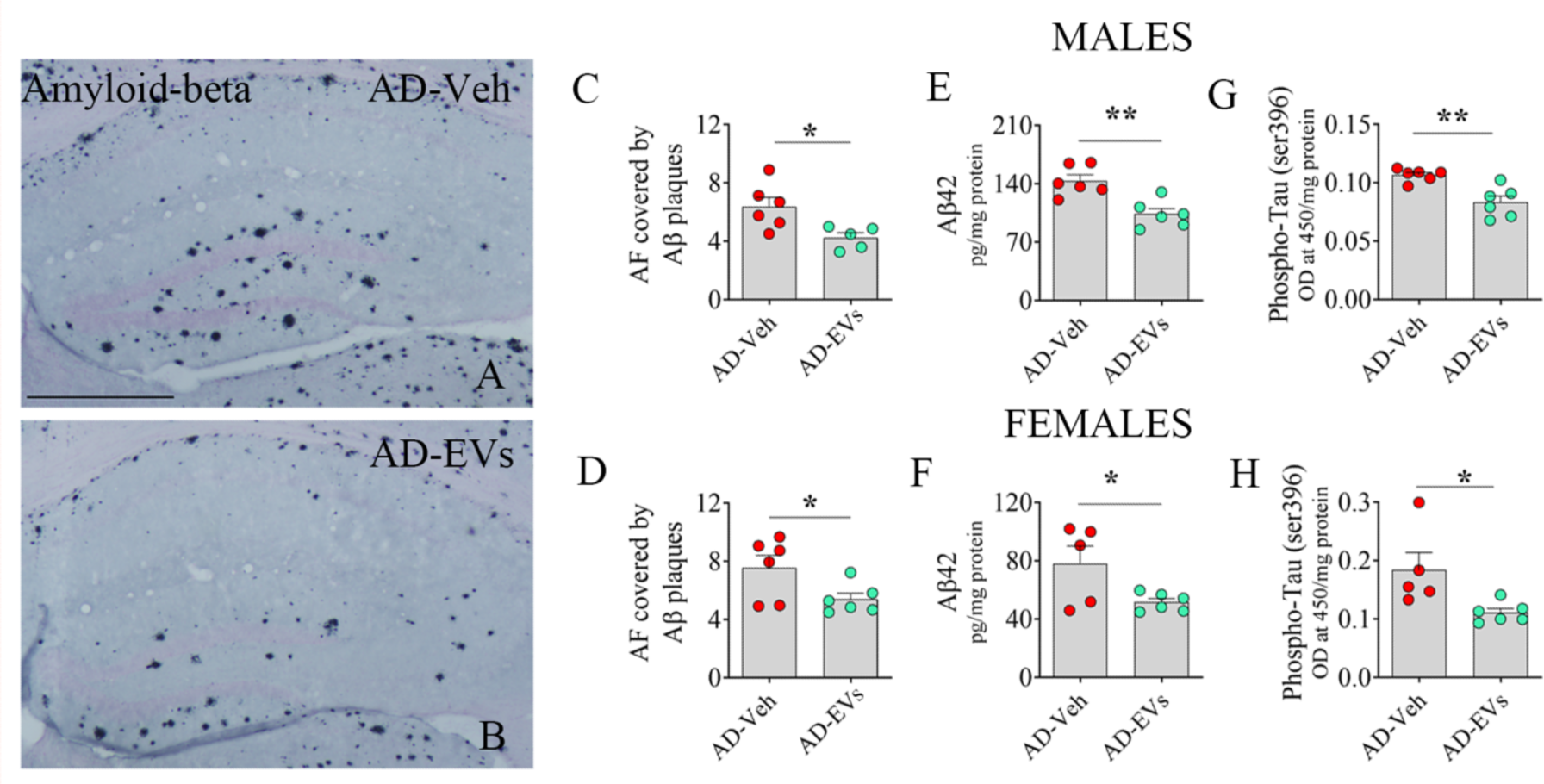
Intranasal administration of extracellular vesicles from human induced pluripotent stem cell-derived neural stem cells (hiPSC-NSC-EVs) to 5xFAD mice reduced amyloid plaques and phosphorylated tau. The bar charts C-H compare the concentration of Aβ42 (E-F) and p-tau (G-H) in males (E, G) and females (F, H) between AD-Veh and AD-EVs groups. Scale bar, A-B =500 μm; *, p<0.05; **, p<0.01; NS, not significant.

