## Supplemental File for "Intranasally Administered EVs from hiPSC-derived NSCs Alter the Transcriptomic Profile of Activated Microglia and Conserve Brain Function in an Alzheimer’s Model"

### A. Methods

#### *A.1. Purification and characterization of extracellular vesicles (EVs) from human induced pluripotent stem cell (hiPSC)-derived neural stem cell (NSC) culture media*

The conditioned media (CM) collected from passage 11 hiPSC-NSC cultures was combined and stored at -20$^o$ C until further use. This media containing EVs was next processed by low-speed centrifugation at 3000 rpm for 10 minutes, followed by filtration through a 0.22 µm filter (GE Healthcare, Uppsala, Sweden) to remove any cellular debris and larger particles. The filtered media was then concentrated 5-7 times using Amicon 30 kDa cut-off ultra-filtration devices (Millipore, Burlington, MA, USA). A chromatography column of 1.5 x 12 cm (Biorad, CA, USA) was filled with 10 mL of Q-Sepharose fast flow from GE Healthcare and equilibrated with 100 mL of equilibration buffer. The concentrated CM was added, followed by a washing step. The EVs were selectively eluted using an elution buffer containing 50 mM Tris and 1000 mM sodium chloride (NaCl) of pH 8.0. Fractions were collected at a flow rate of 1 mL/min, and the elution of EVs was continuously monitored using Nanoparticle tracking analysis (NTA).

After the AEC process, the fractions that contain EVs are pooled and concentrated using an ultrafiltration device with a 50 kDa cut-off. Next, size exclusion chromatography (SEC) was performed on a column composed of 20 mL of Sephacryl S-500 High Resolution (GE Healthcare, Uppsala, Sweden). The EVs are size fractionated using a mobile phase containing 50 mM phosphate buffer and 200 mM NaCl of pH 7.4, and fractions are collected at a 1 mL/min flow rate. The elution of EVs is constantly monitored by quantifying the total protein content using the bicinchoninic acid (BCA) method and tracked by NTA. The fractions that contain the highest number of EVs with minimal protein content are pooled, concentrated, and stored at -20$^o$ C for further use. The EVs obtained from every batch of NSC cultures are subjected to western blot or ELISA to confirm the presence of EV-specific proteins as described elsewhere[1]. Transmission electron microscopy (TEM) of hiPSC-NSC-EVs were performed at the Image Analysis Laboratory, Texas A&M Veterinary Medicine & Biomedical Sciences, as described in our previous report[1].

#### *A.2. Generation of iMicroglia from hiPSCs*

Four-day-old hiPSC cultures were re-seeded with 60,000 cells per $cm^2$ and incubated with E6 medium supplemented with 7.5 ng/mL Activin A, 30 ng/mL bone morphogenetic protein-4 (BMP-4), 3 µM CHIR (a selective glycogen synthase kinase-3 [GSK-3] inhibitor), and 10 µM Rho Kinase (ROCK) inhibitor. After 18 hours, the medium was replaced with fresh E6 medium containing 10 ng/mL Activin A, 40 ng/mL BMP-4, and 2 µM IWP-2 (Wnt processing and secretion inhibitor). The next day, E6 medium with 10 ng/mL Activin A, 40 ng/mL BMP-4, 2 µM IWP-2, and 20 ng/mL fibroblast growth factor-2 (FGF-2) was added. On day 3, cells were dissociated and seeded on a Matrigel plate with a cell density of 50,000 cells per $cm^2$. Vascular endothelial growth factor (VEGF, 15 ng/mL), FGF-2 (5 ng/mL), and ROCK inhibitor (10 µM) were added to the E6 medium. The same fresh medium was added the following day but without ROCK inhibitor. On days 5 and 6, cells were grown in an E6 medium containing 15 ng/mL VEGF, 5 ng/mL FGF-2, and 200 ng/mL stem cell factor (SCF). From day 7 to 9, the E6 medium was used as basal medium supplemented with 100 ng/mL SCF, 30 ng/mL thrombopoietin (TPO), and 30 ng/mL interleukin-3 (IL-3). During this step, floating cells were re-seeded on the culture dish. On day 10, floating and semi-adhered cells were collected and replated on Primaria-coated plates or Poly-D-Lysine pre-coated plates with a cell density of 20,000 cells per $cm^2$. For the microglial cell maturation, RPMI1640 was used as a base medium supplemented with 10% fetal bovine serum (FBS), macrophage-colony stimulating factor (M-CSF, 200 ng/mL), and IL-34 (100 ng/mL). The cell maturation was confirmed by immunostaining using a primary antibody for TMEM119 (BioLegend, San Diego, CA, USA).

#### *A.3. Assessment of a hippocampus-dependent cognitive function using an object Location test (OLT)*

A behavioral task focused on identifying cognitive impairment in animals by observing their preference to explore an object moved to a new location was employed. The ability to recognize minor changes in the immediate environment indicates intact hippocampus-dependent cognitive function. The animals underwent an object location test (OLT) consisting of three trials (T1-T3), each lasting 5 minutes, with a 15-minute inter-trial interval (ITI). The details of the test were described in previous studies[2-4]. In T1, the animal explored an open field apparatus (50 cm W, 50 cm L, 38 cm H) for 5 minutes to get familiar with the environment. In T2, after a 15-minute ITI, the animal spent 5 minutes with two similar objects placed on one side of the chamber. The AnyMaze video tracking system was used to determine if the animal explored both objects in this phase. In T3, the animal spent 5 minutes in the same apparatus with one object moving to a new location. The apparatus was cleaned with 70% alcohol and air-dried after each trial. The AnyMaze video-tracking system recorded the movement of animals in T2-

T3. The animals that explored objects for ≥20 seconds in T2 and ≥8 seconds in T3 were included for data analysis because the ability to identify the object in the novel place in T3 requires the memory for the location of objects in T2, and such aptitude needs exploration of objects for significant periods in T2[5]. The study calculated and compared several results, such as the percentage of time spent exploring the object in a familiar place (OIFP) versus the object in a novel place (OINP) in T3, the total object exploration times (TOETs) in T2, and the discrimination index (DI). The DI values were compared to the hypothetical mean (zero) using a one-sample t-test.

***A.4. Evaluation of pattern separation ability using a pattern separation test (PST)***

The pattern separation function, a cognitive ability that allows encoding of similar but not identical experiences in a non-overlapping way[6-7], was measured through a PST. Maintenance of this function depends upon the integrity of the dentate gyrus of the hippocampus, including the extent of neurogenesis. After acclimatization to the open field apparatus (T1), each mouse successively explored two different sets of identical objects (object types 1 and 2) placed on distinct types of floor patterns (Patterns 1 and 2, P1 and P2) for 5 minutes each in the two acquisition trials T2 and T3 separated by 60 minutes. Sixty minutes later, in the testing phase (T4), each mouse explored an object from T3 (which is now a familiar object, FO) and an object from T2 (which is now a novel object, NO) placed on the floor pattern employed in T3 (P2). The choice to explore the NO on P2 more than the FO on P2 reflects the ability of the animal to distinguish similar but not identical experiences in a non-overlapping fashion. Novel object DI values were calculated and compared to the hypothetical mean (zero) using a one-sample t-test. Only animals that explored objects for >20 seconds in T2-T3 and ≥8 seconds in T4 were included for data analysis.

***A.5. Appraisal of Anhedonia using a sucrose preference test (SPT)***

Alzheimer's disease (AD) is also associated with symptoms of depressive-like behavior or anhedonia. To assess the extent of anhedonia in vehicle or hiPSC-NSC-EVs treated 5xFAD mice, we used an SPT, as described in a previous report[8]. In this test, anhedonia is indicated by a reduced preference for sweet fluids such as sucrose- or saccharin-containing water. The test involved monitoring animals for four consecutive days. On Day 1, mice were housed individually and given free access to two identical bottles containing 1% sucrose solution. They were also allowed to eat food as much as they wanted. On

Day 2, one of the bottles was replaced with a new bottle containing regular drinking water, and the animals were again given ad libitum access to food. On Day 3, the animals were deprived of water and food for 22 hours. On Day 4, the animals' preference for consuming sucrose-containing water over regular water was tested by giving them access to two bottles for 2 hours, one containing 1% sucrose solution and the other containing regular water. The volume of the bottles was measured, and the mice were placed back in their previous housing conditions with ad libitum access to water and food. Anhedonia was assessed by comparing the sucrose preference rate (SPR) between study groups. In addition, the preference for sucrose-containing water over regular water within individual study groups was also measured.

#### *A.6. Preparation of brain tissues for immunohistochemical and biochemical analyses*

For immunohistochemical studies, subgroups of animals from all groups were anesthetized with isoflurane, and then transcardial perfusion with 4% paraformaldehyde was performed. The entire brain was removed from each animal and post-fixed in 4% paraformaldehyde at 4°C. After treatment with a gradient of sucrose solutions, thirty-micrometer thick sections were cut coronally from the entire forebrain using a cryostat and stored at -20 degrees Celsius in an antifreeze solution until further use. For biochemical studies, following deep anesthesia, fresh brain tissues were harvested from additional subgroups of animals and snap-frozen using liquid nitrogen and stored at -80°C until needed. Tissues were thawed to room temperature and micro-dissected to remove the hippocampus tissue. Hippocampal tissue lysates were prepared using tissue extraction buffer (ThermoFisher Scientific, Waltham, Massachusetts, USA).

#### *A.7. Immunohistochemical staining for visualizing amyloid-beta and IBA-1*

The process for preparing the tissue for immunohistochemical analysis is described in our previously published studies[1,9-14]. In summary, the tissue sections were first treated with PBS comprising 10% normal goat serum (NGS) and 0.1% Triton X-100 for 30 minutes at room temperature. The sections were then incubated overnight at $4^{O}$ C in the respective primary antibody solution: rabbit anti-amyloid beta (1:500, Invitrogen) or rabbit anti-IBA-1 (1:1000-Abcam) solution in PBS containing 3% serum and 0.1% Triton X-100. Afterward, the tissues were incubated in a biotinylated goat anti-rabbit IgG solution (1:200) containing 3% NGS for one hour at room temperature, followed by incubation in the avidinbiotin complex (ABC) reagent for 1 hour. These tissue sections were then developed in Vector SG solution for 6-8 minutes. They were washed in distilled water, mounted, and dehydrated with graded ethyl alcohol, and the cover slipped with permount.

#### *A.8. Single, dual, or triple immunofluorescence methods*

Single, dual, or triple immunofluorescence procedures were employed to visualize the following markers. 1) Microglia containing PKH26+ hiPSC-NSC-EVs. 2) Microglia positive for IBA-1 and CD68. 3) IBA-1 positive microglia containing complexes positive for nucleotide-binding domain leucine-rich repeat and pyrin domain-containing receptor 3 (NLRP3) and apoptosis-associated speck-like protein containing a CARD (ASC). The sections were washed in PBS, blocked with 10% normal donkey serum (NDS), and incubated overnight at $4^{O}$ C in individual primary antibodies (for single immunofluorescence) or a cocktail of two or three primary antibodies (for dual and triple immunofluorescence) in NDS. The sections were incubated the following day with matching secondary antibodies for 60 minutes. Then, the sections were rinsed in PBS and coverslipped with a slow fade/antifade mounting medium (Invitrogen). The primary antibodies comprised goat anti-IBA-1 (1:1000, Abcam), rabbit anti-IBA-1 (1:1000, Abcam), goat anti-NLRP3 (1:500, Millipore), and mouse anti-ASC (1:500, Santa Cruz). The secondary antibodies used were donkey anti-mouse IgG-Alexa Flour 594 (1:200, Invitrogen), donkey anti-goat IgG-Alexa Fluor 488 (1:200, Invitrogen), donkey anti-rabbit IgG-Alexa Fluor 405 (1:200, Invitrogen).

#### *A.9. Preparation of brain tissue lysates for biochemical studies*

Hippocampal tissues from all groups were micro-dissected. The tissues were lysed through sonication in a tissue extraction reagent (Invitrogen) containing protease inhibitor (1:100 dilution, Sigma) for 15-20 seconds at $4^{O}$ C. The resulting solution was centrifuged at $4^{o}$C for 10 minutes (15000g), and the supernatant was aliquoted and stored at $-80^{O}$ C until further use. The lysates were used to measure the concentration of multiple markers.

#### *A.10. Enzyme-linked immunosorbent assays (ELISAs) for measuring various proteins*

The lysates from the hippocampus were dispensed as recommended to 96-well plates that were pre-coated with specific capture primary antibodies and then incubated with gentle shaking. After several rinses with a wash buffer, specific biotinylated detection antibodies were added and incubated as the manufacturer's protocol recommended. The plates were washed, incubated in a streptavidin-HRP conjugate solution, and rewashed. The plates were then treated sequentially with detection and stop reagents and read at 450 nm. The concentration of each protein was measured using standard graphs. The protein concentration in the lysate was normalized to mg of protein for the hippocampus lysate.

### B. Results

#### *B.1. Characterization of hiPSC-NSC-EVs*

NanoSight analysis of EVs isolated from P11 hiPSC-NSC cultures through AEC and SEC methods revealed the presence of EVs with a mean size of ~145 nm (Supp Fig. 1 [A]). Western blots validated that hiPSC-NSC-EVs expressed multiple EV-specific markers such as CD63, CD81, and ALIX. Also, the EV lysates lacked the expression of deep cellular proteins such as calnexin and cytochrome C (Supp Fig. 1 [B]). Furthermore, TEM analysis confirmed the size and classic morphology of EVs enclosed by double membranes (Supp Fig. 2 [C]).

#### *B.2. Supplement Figure 1*

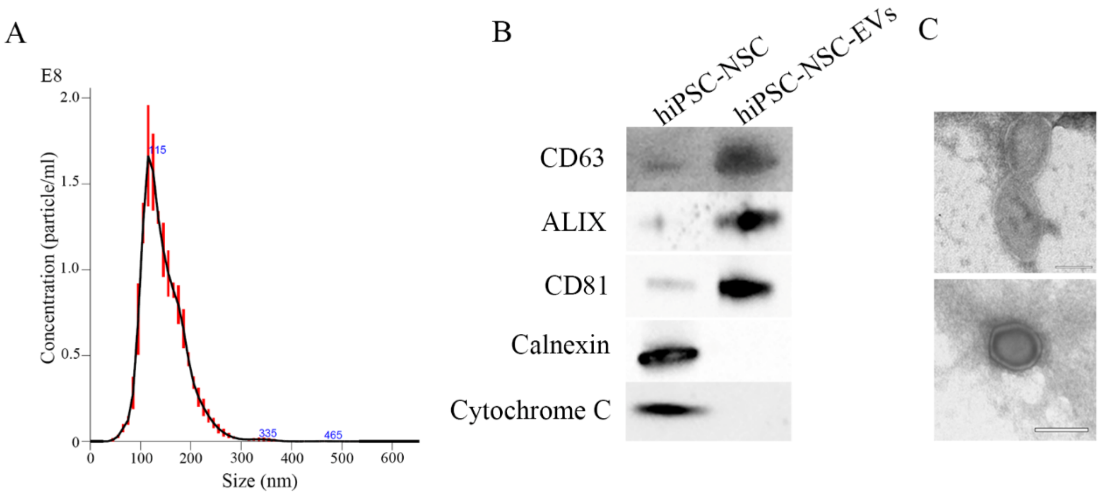

***Supplement Figure 1:*** *Characterization of hiPSC-NSC-EVs. Graph A shows the size and number of hiPSC-NSC-EVs measured with a NanoSight. The blots in B demonstrate EV-specific proteins CD63, CD81, and ALIX and the absence of the deep cellular proteins, cytochrome c and calnexin, in hiPSC-NSC-EVs. The images in C display the morphology and size of hiPSC-NSC-EVs visualized through transmission electron microscopy: scale bar, 100 nm.*

***B.3. Supplement Figure 2***

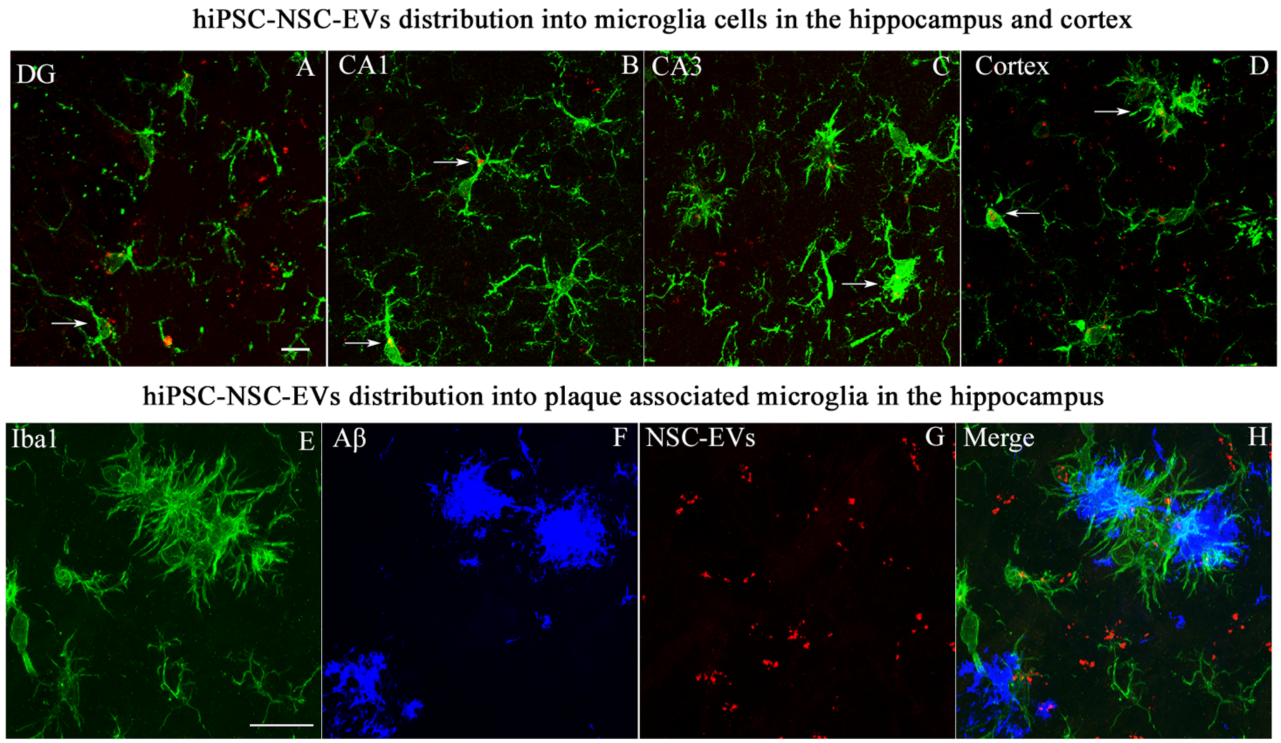


***Supplemental Figure 2:*** *Intranasally administered extracellular vesicles from human induced pluripotent stem cell-derived neural stem cells (hiPSC-NSC-EVs) incorporated into IBA-1+ microglia in the hippocampus and the cerebral cortex of 5xFAD mouse. Figures A-D illustrate the incorporation of EVs into Iba1 + microglia in different brain regions of 5xFAD mice (A: DG; B: CA1; C: CA3; D: Cortex) at 45 min post-administration. Figures E-H show the incorporation of EVs into IBA-1+ plaque-associated microglia; Scale bar, A-H=10 µm.*

***B.4. Supplement Figure 3***

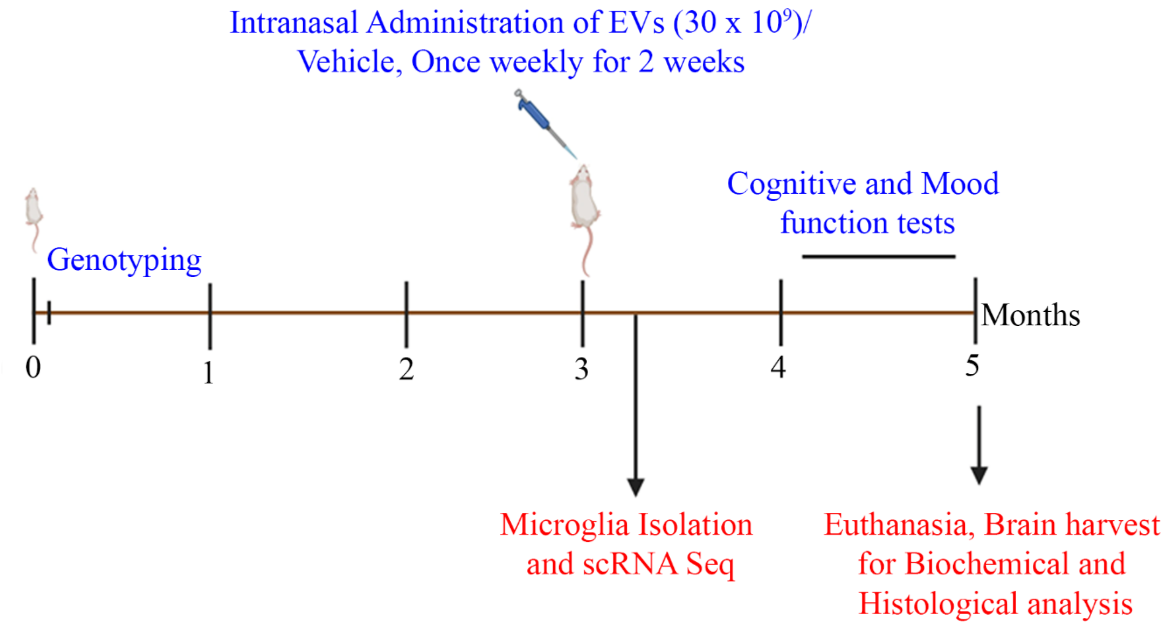


***Supplement Figure 3:*** *The experimental design showing the time points of hiPSC-NSC-EVs treatment, cognitive and mood function tests, euthanasia, and brain tissue harvest.*

*B.4. Supplement Figure 4*

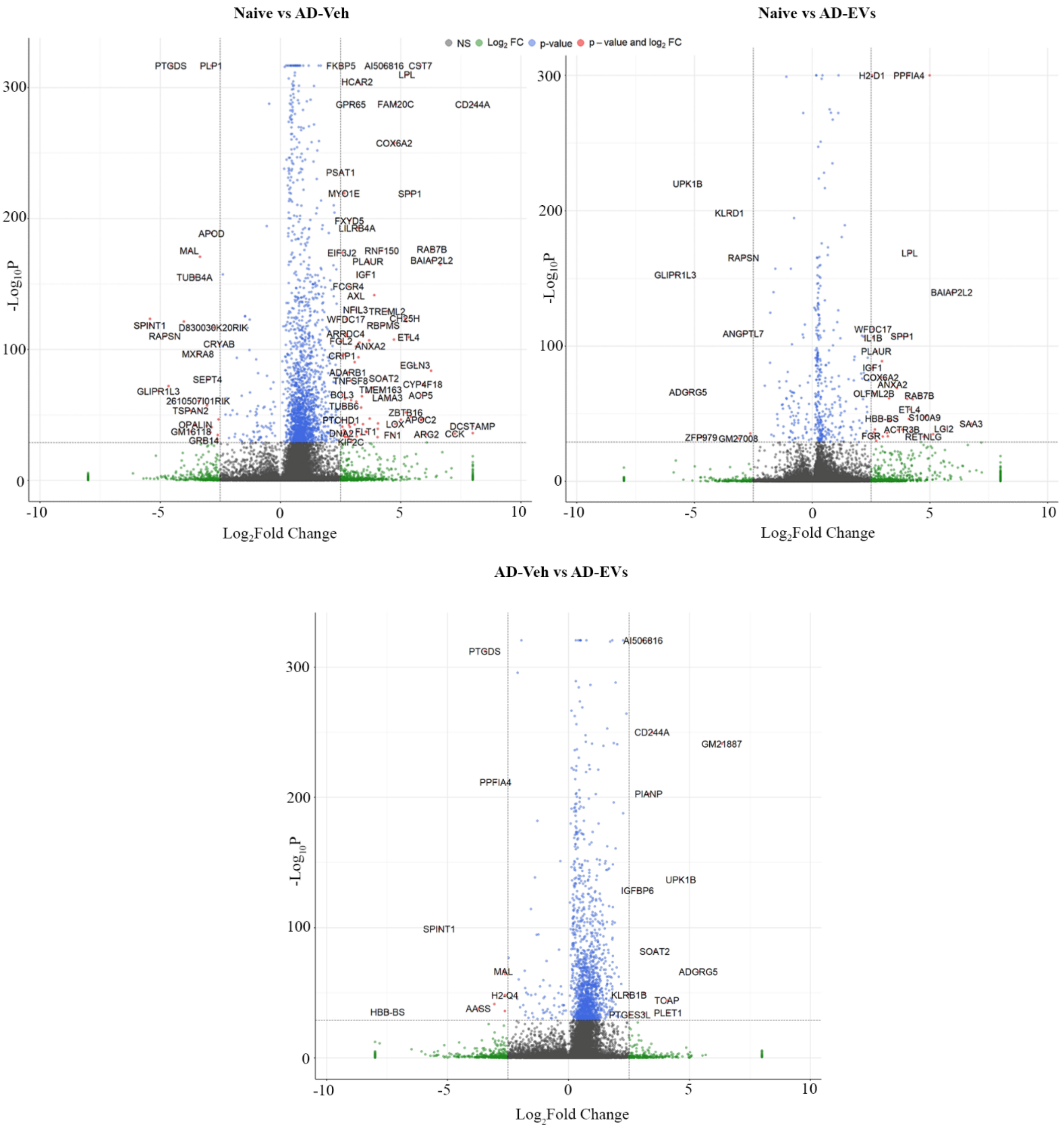


*Supplement Figure 4: The figures show volcano plots of differential gene expression in microglia between naïve vs AD-Veh, naïve vs AD-EVs and AD-Veh vs AD-EVs groups, measured through single-cell RNA sequencing study.*

#### *B.5. Supplement Figure 5*

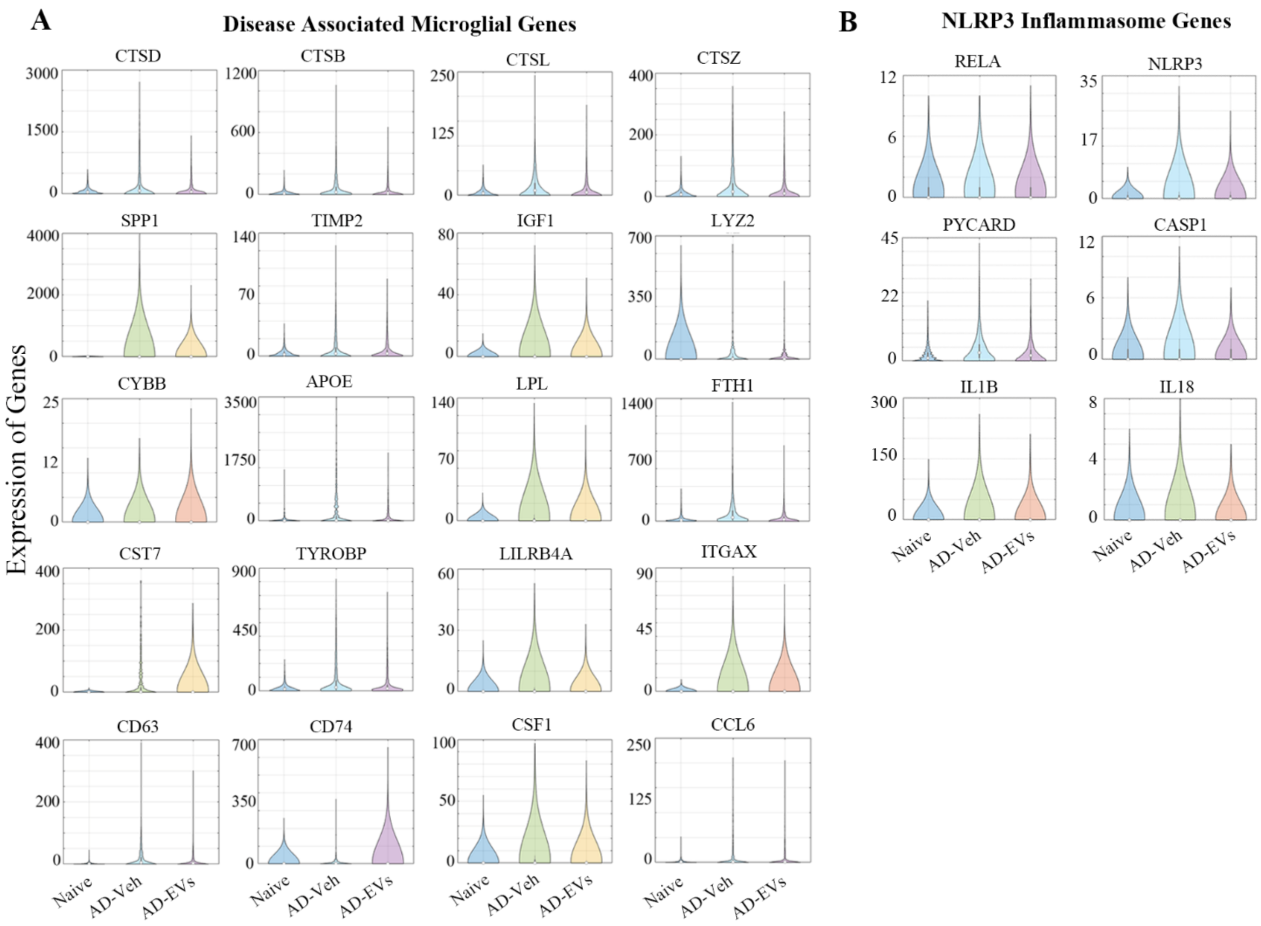


***Supplement Figure 5**: The violin plots compare the expression of various genes linked to disease-associated microglia (DAM; A) and* NOD-, LRR- and pyrin domain-containing protein 3 (*NLRP3; B) inflammasomes.*

#### B.6. Supplement Figure 6

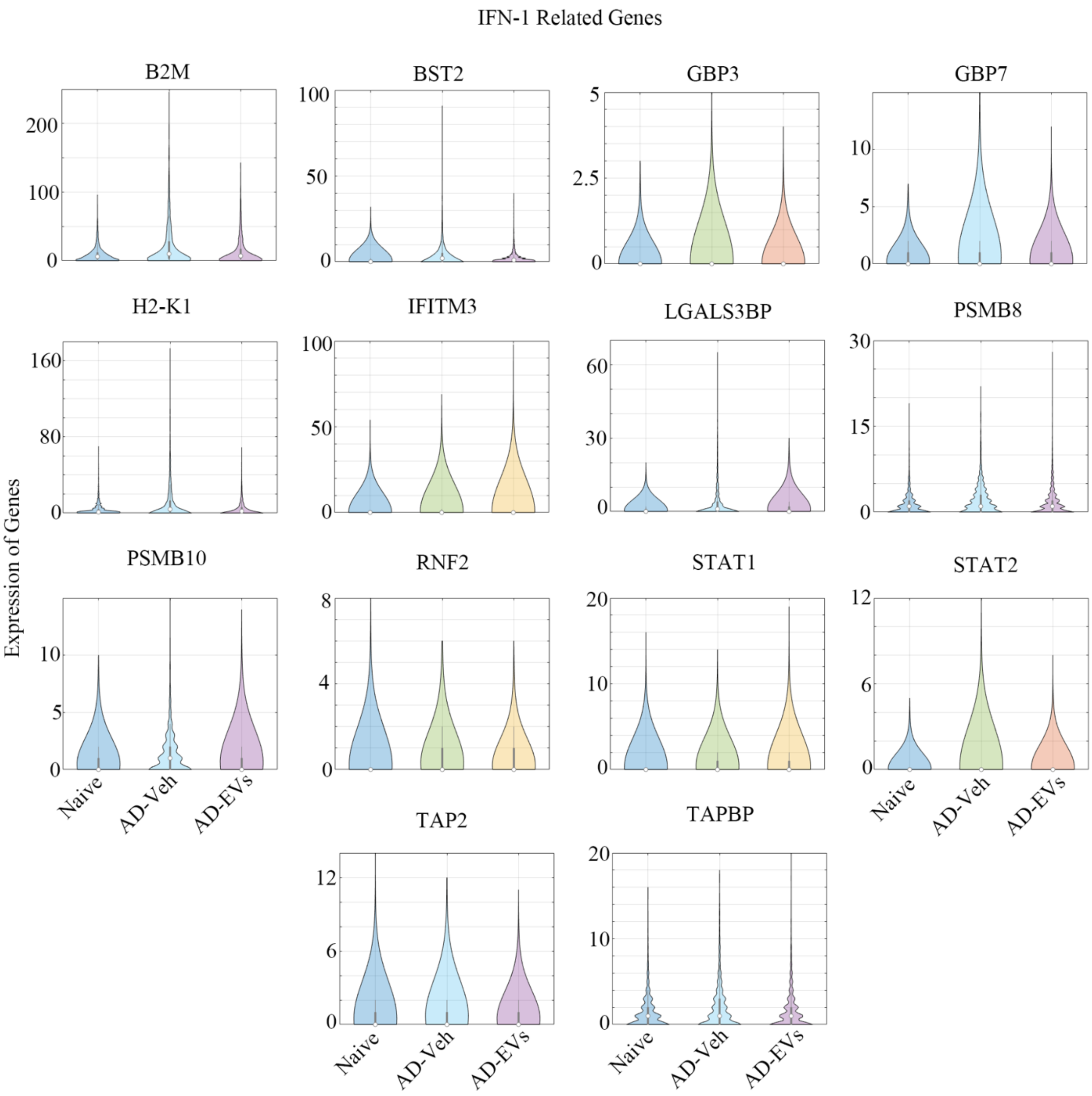


***Supplement Figure 6****: The violin plots compare the expression of various genes linked to Interferon 1 (IFN 1) signaling between naïve, AD-Veh, and AD-EVs groups.*

***B.7. Supplement Figure 7***

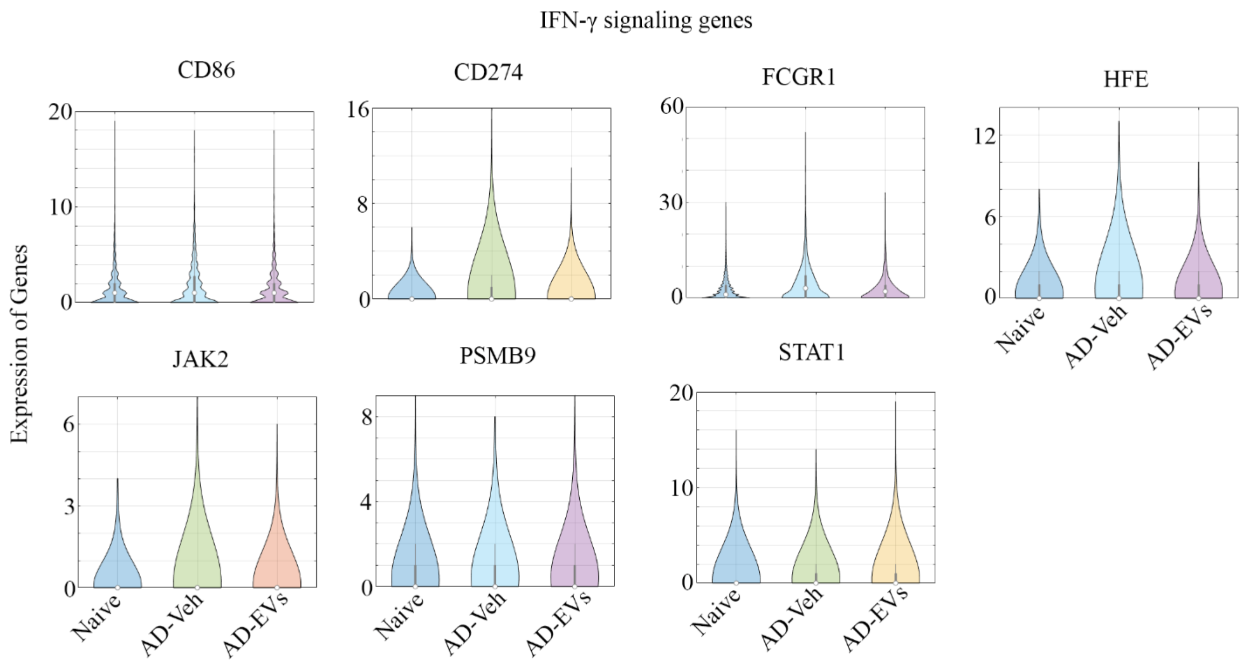


***Supplement Figure 7**: The violin plots compare the expression of various genes linked to IFN-γ signaling between naïve, AD-Veh, and AD-EVs groups.*

***B.8. Supplement Figure 8***

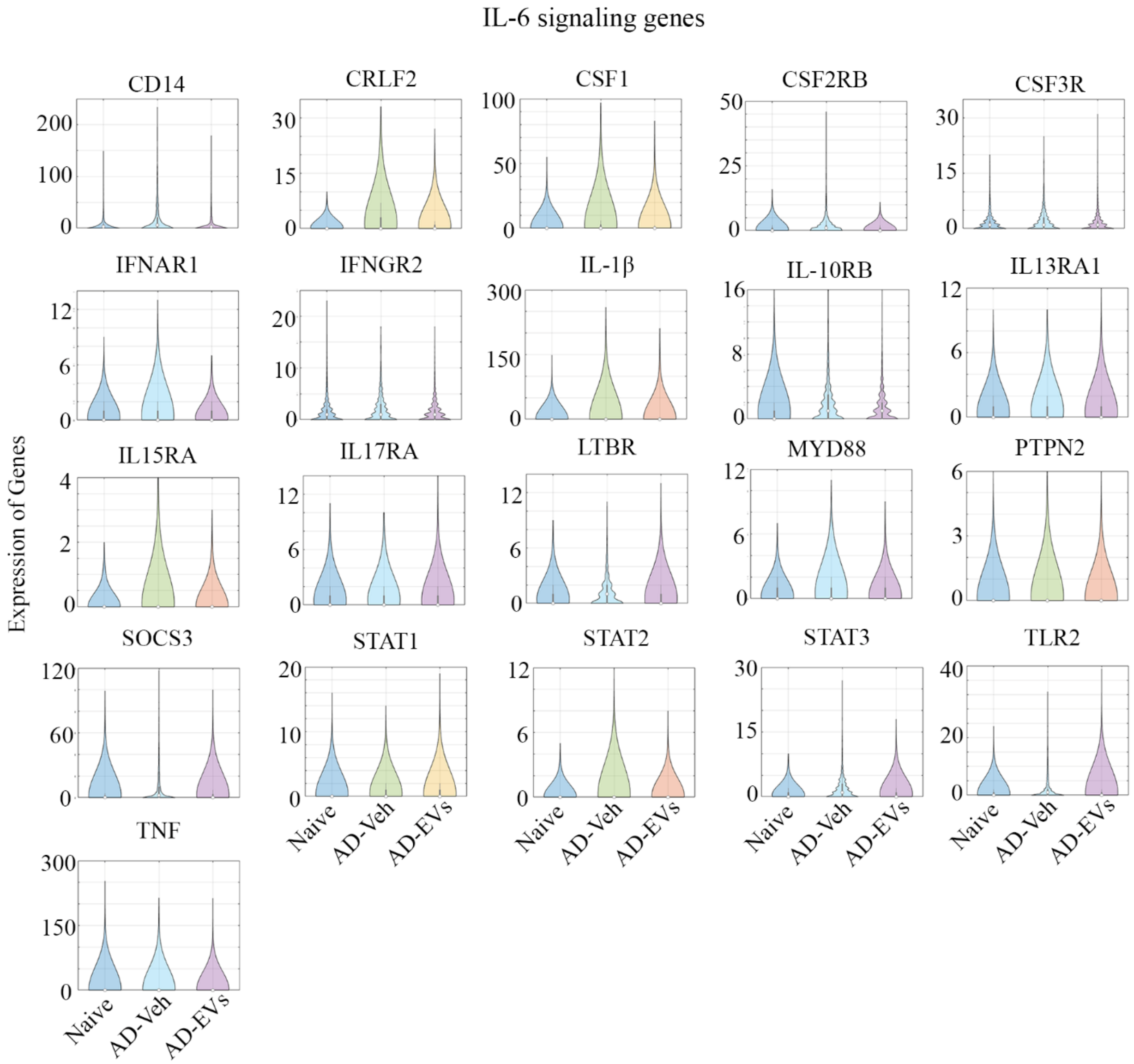


***Supplement Figure 8****: The violin plots compare the expression of various genes linked to IL-6 signaling between naïve, AD-Veh, and AD-EVs groups.*

***B.9. Supplement Figure 9***

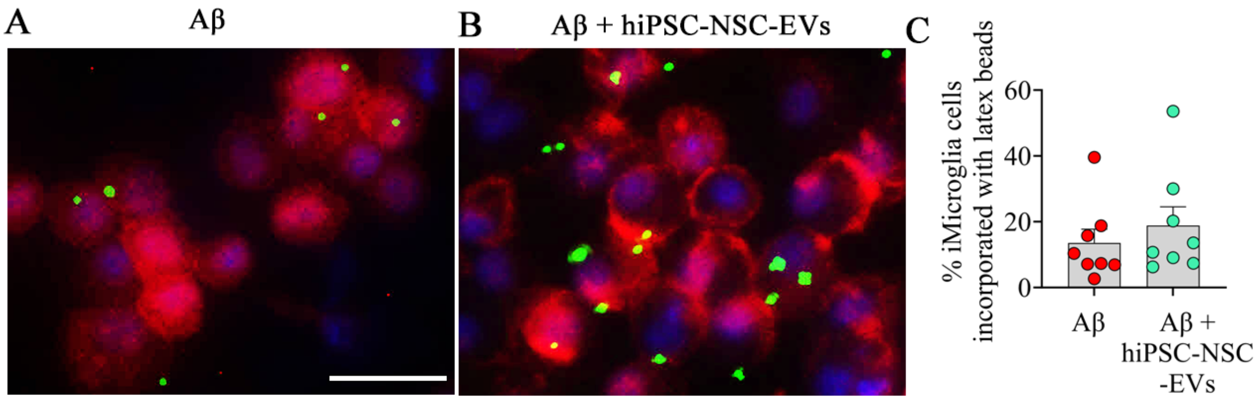


***Supplement Figure 9:*** *The images show the phagocytosis of latex beads by iMicroglia exposed to Aβ oligomers alone (A) or Aβ oligomers and hiPSC-NSC-EVs (B). The bar chart (C) compares percentages of iMicroglia incorporating latex beads between iMicroglia cultures exposed to Aβ oligomers alone or Aβ oligomers and hiPSC-NSC-EVs; Scale bar 100 µm.*

***B.10. Supplement Figure 10***

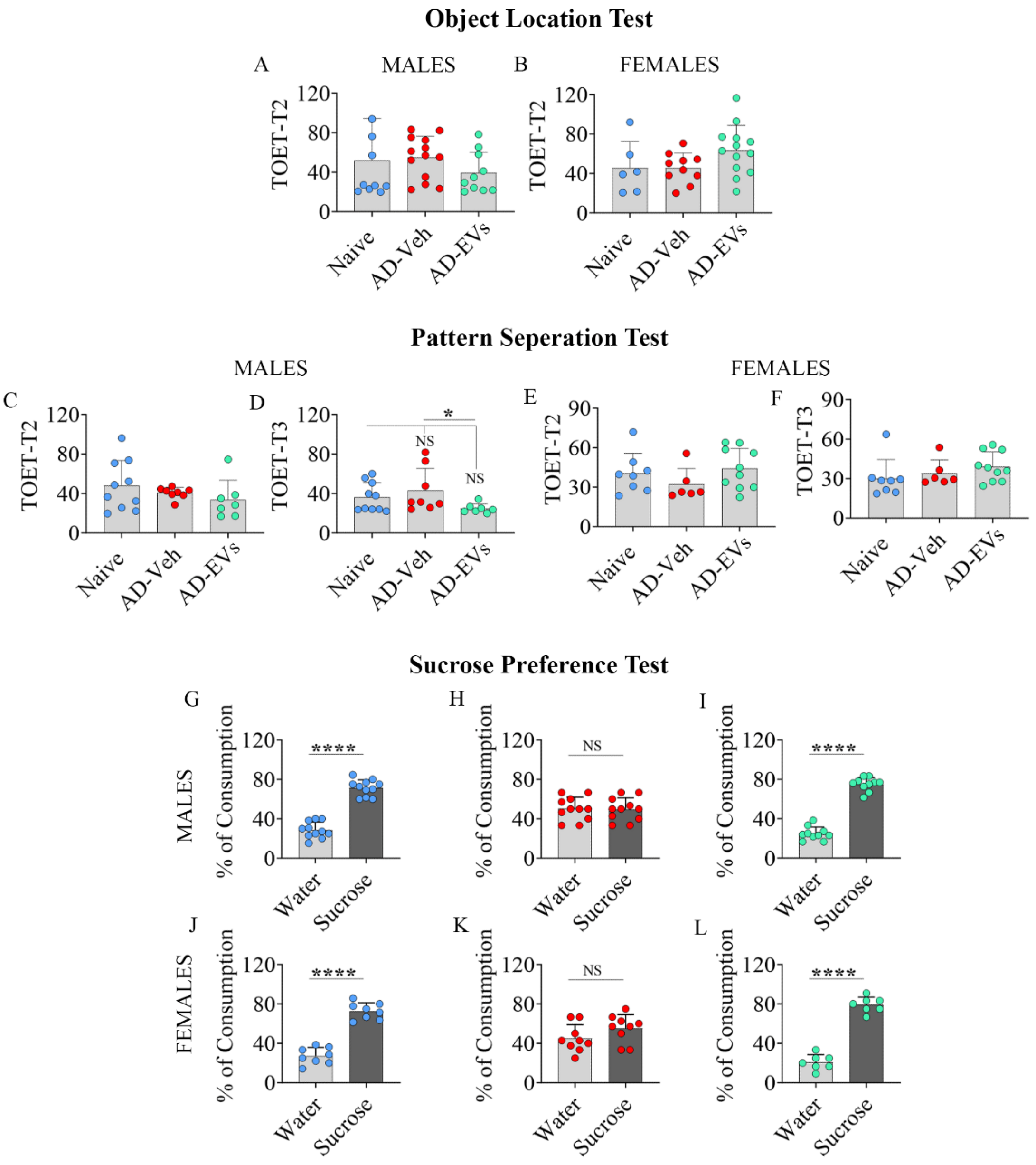


***Supplementary Figure 10:*** *Comparison of total object exploration times (TOETs) in the object location test (OLT) and the pattern separation test (PST), and percentages of fluid consumption in the sucrose preference test (SPT). The bar charts A-B compare TOETs in males (A) and females (B) from trial 2 (T2) of an OLT between naive, AD-Veh, and AD-EVs groups. The bar charts in C-F compare TOETs in males (C, D) and females (E, F) from T2 (C, E) and T3 (D, F) of a PST between naive, AD-Veh, and AD-EVs groups. The bar charts G-L compare standard water and sucrose-containing water consumption percentages in males (G-I) and females (J-L) of naive (G, J), AD-Veh (H, K), and AD-EVs (I, L) groups. *, p<0.05; ****, p<0.0001; NS, not significant.*
